# The variability of multidimensional diffusion-relaxation MRI estimates in the human brain

**DOI:** 10.1101/2024.05.20.594998

**Authors:** Eppu Manninen, Shunxing Bao, Bennett A. Landman, Yihong Yang, Daniel Topgaard, Dan Benjamini

## Abstract

Diffusion-relaxation correlation multidimensional MRI (MD-MRI) replaces voxel-averaged diffusion tensor quantities and R_1_ and R_2_ relaxation rates with their multidimensional distributions, enabling the selective extraction and mapping of specific diffusion-relaxation spectral ranges that correspond to different cellular features. This approach has the potential of achieving high sensitivity and specificity in detecting subtle changes that would otherwise be averaged out. Here, the whole brain characterization of MD-MRI distributions and derived parameters is presented and the intrascanner test–retest reliability, repeatability, and reproducibility are evaluated to promote the further development of these quantities as neuroimaging biomarkers. We compared white matter tracts and cortical and subcortical gray matter regions, revealing notable variations in their diffusion-relaxation profiles, indicative of unique microscopic morphological characteristics. We found that the reliability and repeatability of MD-MRI-derived diffusion and relaxation mean parameters were comparable to values expected in conventional diffusion tensor imaging and relaxometry studies. Importantly, the estimated signal fractions of intra-voxel spectral components in the MD-MRI distribution, corresponding to white matter, gray matter, and cerebrospinal fluid, were found to be reproducible. This underscores the viability of employing a spectral analysis approach to MD-MRI data. Our results show that a clinically feasible MD-MRI protocol can reliably deliver information of the rich structural and chemical variety that exists within each imaging voxel, creating potential for new MRI biomarkers with enhanced sensitivity and specificity.

## 1 Introduction

Diffusion-relaxation multidimensional MRI (MD-MRI) acquisition integrates meso- and microstructural probes (Callaghan, 2011) with chemical composition sensitivity (Tofts, 2003). While voxel average relaxation and diffusion metrics have the capacity to be transformed into basic estimates of, e.g., myelin fraction (Mackay et al., 1994; Vasilescu et al., 1978) or general orientation of nerve fiber bundles (Basser et al., 1994) within individual millimeter-scale image voxels, achieving a more precise interpretation is hindered by the complex reality that each voxel comprises numerous cells with varying sizes, shapes, and orientations (Livet et al., 2007). This heterogeneity creates considerable uncertainty when establishing connections between the observable metrics and the specific microscopic details we are often interested to discern.

While the classical diffusion tensor imaging (DTI) model (Basser et al., 1994) has proven sensitive to probing brain microstructure in a range of scenarios and applications (Soares et al., 2013), it is encumbered by inherent limitations that impede its applicability and the precision of its interpretations (Novikov et al., 2018). To enhance specificity, various multicomponent biophysical models have been developed and successfully applied to study brain white matter (WM) (Assaf & Basser, 2005; Benjamini et al., 2016; Stanisz et al., 1997; Veraart et al., 2020; Zhang et al., 2012). While these models offer advantages in specificity and interpretability, they rely on strict assumptions and incorporate fixed parameters to reduce fitting ambiguities (Novikov et al., 2018), which may not be valid in disease states or even in heterogeneous gray matter (GM) regions (Jespersen et al., 2019; Lampinen et al., 2019; Novikov et al., 2018; Veraart et al., 2019).

An alternative approach is to simultaneously encode diffusion-relaxation data and apply a model-free estimation of distributions of quantitative metrics. These distributions provide information regarding the amounts of individual diffusion and relaxation properties of distinct water populations, and their correlations (Hürlimann & Venkataramanan, 2002; Silva et al., 2002). The obtained information has been shown invaluable to the study of tissue microstructure and brain connectivity (de Almeida Martins et al., 2021; de Almeida Martins et al., 2020; Kundu et al., 2023; Peled et al., 1999; Reymbaut et al., 2021; Stanisz & Henkelman, 1998; Yon et al., 2020) as well as pathology (Benjamini et al., 2020; Benjamini et al., 2022; Kim et al., 2017; Slator et al., 2021; Wei et al., 2022). The employment of diffusion acquisition strategies that incorporate free gradient waveforms (Sjölund et al., 2015) facilitates the exploration of both the frequency-dependent and tensor-related characteristics of the diffusion encoding spectrum **b**(ω) (Lasič et al., 2022; Lundell & Lasič, 2020). This approach, usually termed multidimensional (MD) MRI, allows for the examination of diffusion-relaxation correlation measures that vary with frequency and time within a unified framework (Narvaez et al., 2022).

An efficient and sparse *in vivo* frequency-dependent MD-MRI acquisition protocol that provides whole brain coverage at 2-mm isotropic resolution was recently introduced (Johnson et al., 2024). This proof-of-concept study tested the feasibility of estimating voxelwise distributions of frequency-dependent diffusion tensors, **D**(ω), and longitudinal and transverse relaxation rates, R_1_ and R_2_, from this sparsely acquired MD-MRI data. The rich information contained within the high-dimensional **D**(ω)-R_1_-R_2_ distributions can be distilled into first (means) and second (variances and covariances) moment statistics of, e.g., frequency-dependent isotropic diffusivity and diffusion anisotropy, R_1_, and R_2_, in each voxel. Further, having voxelwise distributions provides access to distinct water populations within a voxel, and therefore enables improved specificity towards WM, GM, and cerebrospinal fluid (CSF) (de Almeida Martins et al., 2020). In turn, this unique intra-voxel information allows to map microstructure-specific diffusion and relaxation properties without relying on biophysical assumptions or models.

In recent studies, the specificity of estimated intra-voxel components using MD-MRI was evaluated in both physical (Benjamini & Basser, 2016; de Almeida Martins & Topgaard, 2016, 2018) and synthetic (Reymbaut et al., 2020) phantoms, and via comparisons with microscopy of fixed tissue samples from the rat and ferret brain (Benjamini & Basser, 2017; Narvaez et al., 2022). The applicability of this framework for *in vivo* human MRI was confirmed by comparing MD-MRI-derived values in the human brain with those reported in the literature (de Almeida Martins et al., 2021; Martin et al., 2021; Naranjo et al., 2021). In this context, our aims are to present a characterization of **D**(ω)-R_1_-R_2_ estimates throughout the human brain and to evaluate their intra-scanner test–retest reliability and repeatability. These efforts contribute to the ongoing development of the MD-MRI framework as a neuroimaging modality.

## 2 Methods

### 2.1 Participants

Ten healthy participants (mean age 48, standard deviation 14.4 years; 4 women) were each scanned twice, a few weeks apart (i.e., a total of 20 scans). Participants were systematically drawn from ongoing healthy cohorts of the National Institute on Drug Abuse (NIDA). Experimental procedures were performed in compliance with our local Institutional Review Board, and participants provided written informed consent. Prior to each scan, NIDA clinical and nursing units conducted COVID-19 testing, urine drug tests, a physical exam, and a questionnaire on pre-existing conditions and daily habits. Exclusion criteria included major medical illness or current medication use, a history of neurological or psychiatric disorders or substance abuse.

### 2.2 Data acquisition

Data were acquired using a 3T scanner (MAGNETOM Prisma, Siemens Healthcare AG, Erlangen, Germany) with a 32-channel head coil. MD-MRI data were acquired with 2-mm isotropic voxel size using a single-shot spin-echo EPI sequence modified for tensor-valued diffusion encoding with free gradient waveforms (Martin et al., 2021; Wetscherek et al., 2015). The acquisition parameters were set as follows: FOV = 228 × 228 × 110 mm^3^, voxel size = 2 × 2 × 2 mm^3^, acquisition bandwidth = 1512 Hz/pixel, in-plane acceleration factor 2 using GRAPPA reconstruction with 24 reference lines, effective echo spacing of 0.8 ms, phase-partial Fourier factor of 0.75, and axial slice orientation.

The acquisition protocol was designed based on previously described heuristic principles (Martin et al., 2021) used to achieve an efficient and sparse MD-MRI acquisition (Johnson et al., 2024). Briefly, in addition to a b=0 ms/μm^2^ volume, numerically optimized (Sjölund et al., 2015) linear, planar, and spherical b-tensors were employed with **b**-tensor magnitude, b, ranging between 0.1 and 3 ms/μm^2^, and **b**-tensor normalized anisotropy, b_Δ_, of –0.5 (planar), 0 (spherical), and 1 (linear). The spectral content of the employed diffusion gradient waveforms was in the range of 6.6 - 21 Hz centroid frequencies, *ω*_*cent*_/2*π*. We note that each **b**-tensor contains a range of frequencies and that the centroid frequency is used only as its representative value. All data inversion and metric calculations use frequency as such and do not rely on reducing the spectral content of **b**(*ω*) to a single number. The datasets were acquired with a single phase encoding direction (anterior to posterior, AP), and an additional *b =* 0 ms/μm^2^ volume with reversed phase encoding direction (PA). Sensitivity to *R*_1_ and *R*_2_ was achieved by acquiring data with different combinations of repetition times, TR = (0.62, 1.75, 3.5, 5, 7, 7.6) s and echo times, TE = (40, 63, 83, 150) ms. The number of concatenations and preparation scans was increased to allow values of TR below 5 s. A total of 139 individual measurements were recorded over 40 min. More details of the acquisition protocol are available in (Johnson et al., 2024). In addition to MD-MRI, fat-suppressed T1W MPRAGE images with TE=3.42 ms, TR=1900 ms, and 1 mm isotropic voxel size were also acquired as structural images. The structural data were utilized in image registrations and region of interest (ROI) segmentations.

### 2.3 MD-MRI data preprocessing

We followed recent recommendations for denoising and preprocessing of sparse MD-MRI data (Johnson et al., 2024). In short, all data volumes were first concatenated and underwent denoising with the MPPCA technique (Veraart et al., 2016). Then, Gibbs ringing correction (Kellner et al., 2016) for partial k-space acquisitions (Lee et al., 2021) was performed, followed by motion and eddy currents distortion corrections using TORTOISE’s DIFFPREP (Rohde et al., 2004) with a physically-based parsimonious quadratic transformation model and a normalized mutual information metric. For susceptibility distortion correction, a T_1_-weighted image was initially converted into a T_2_-weighted image with b=0 s/mm^2^ contrast (Schilling et al., 2019), which was fed into the DR-BUDDI software (Irfanoglu et al., 2015) for AP-PA distortion correction. The final preprocessed data was output with a single interpolation in the space of an anatomical image at native in-plane voxel size.

### 2.4 Estimation of MD-MRI parameters

Solving the joint **D**(ω)-R_1_-R_2_ probability density distribution from the MD-MRI measurements is an ill-conditioned problem. To solve that inversion problem, the estimation of the MD-MRI parameters used in this study followed the procedures described in (Johnson et al., 2024). The multidimensional diffusion MRI toolbox for MATLAB (Nilsson et al., 2018) was used to perform a Monte Carlo inversion (Narvaez et al., 2022) for the preprocessed MD-MRI data. In the inversion, the MD-MRI signal *S*(**b**(*ω*), TE, TR) is represented as a weighted sum of discrete components *c* of the joint probability density distribution **D**(ω)-R_1_-R_2_:

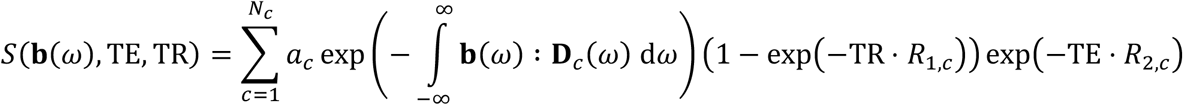

where *N*_*c*_ is the number of fitted components, *a*_*c*_ is the weight of the estimated component *c*, : denotes a generalized dot product, and **D**(*ω*) is an axially symmetric diffusion tensor that depends on the angular frequency of the diffusion gradient waveform. The diffusion tensors are approximated as axisymmetric Lorentzians parameterized by the zero-frequency axial and radial diffusivities [D_||_, D_⊥_], azimuthal and polar angles [*θ, ϕ*], high frequency isotropic diffusivity, D_0_, axial and radial transition frequencies, [Γ_||_, Γ_⊥_], along with longitudinal and transversal relaxation rates [R_1_, R_2_]. In this study, these parameters were sampled in the ranges 0.05<D_||,⊥,0_<5 μm^2^/ms, 0<*θ*<*π*, 0<*ϕ*<2*π*, 0.2<R_1_<2 s^-1^, 1<R_2_<30 s^-1^, and 0.01<Γ _||/⊥_<10000 s^-1^. The Monte Carlo inversion used non-negative least squares with quasi-genetic filtering, bootstrapping, and constraints for the MD-MRI parameter estimates (Benjamini, 2020; de Almeida Martins & Topgaard, 2018). As a result, for each imaging voxel, 300 solutions (bootstraps) of the **D**(ω)-R_1_-R_2_ probability density distribution with up to 10 weighted components (for each bootstrap) were estimated. Voxel-wise **D**(*ω*)-R_1_-R_2_ distributions in the primary analysis space [D_||_, D_⊥_, *θ, ϕ*, D_0_, Γ_||_, Γ_⊥_, R_1_, R_2_] were evaluated at selected values of ω⁄2*π* within the narrow 6.6 - 21 Hz experimental window, giving a set of *ω*-dependent distributions in the [D_||_(*ω*), D_⊥_(*ω*), *θ, ϕ*, R_1_, R_2_] space. It should be noted that the highest frequencies reached for b-values of 0.5, 1.5, and 3 ms/μm^2^ were 21, 15, and 11 Hz, respectively. We further chose to express the axial and radial diffusivities as isotropic diffusivity, D_iso_, and squared normalized anisotropy *D*_Δ_ ^2^ (Conturo et al., 1996).

Two-dimensional projections of the full 6D **D**(*ω*)-R_1_-R_2_ distributions were constructed to facilitate visualization and to allow investigation of certain ROIs. We visualized 2D distributions by projecting and mapping the weights of the discrete components from all bootstrap iterations onto 64 × 64 meshes in the 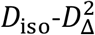, *D*_iso_-*R*_1_, and *D*_iso_-*R*_2_ planes. These voxel-wise 2D projections were then summed over entire ROIs within each scan, normalized, and finally averaged across all 20 scans to produce characteristic distributions.

The estimated MD-MRI parameters assessed for reliability and repeatability in this study included the expectation E[*x*] (mean) values of isotropic diffusivity *D*_iso_ (E[*D*_iso_]), squared diffusion anisotropy 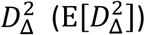, *R*_1_ relaxation rate (E[*R*_1_]), and *R*_2_ relaxation rate (E[*R*_2_]). The effects of restricted diffusion were quantified by a finite difference approximation of the rate of change of the diffusivity metrics with frequency ω⁄2*π* within the investigated window (6.6 to 21 Hz), expressed as Δ⁄2*π* [*x*] of *D*_iso_ and 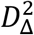 statistics (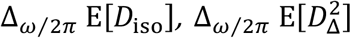, respectively) (Johnson et al., 2024; Narvaez et al., 2022). The two-dimensional diffusivity—diffusion anisotropy space can be used to delineate sub-voxel components corresponding to different microstructural environments (Pierpaoli et al., 1996; Topgaard, 2019). Sub-voxel components were expressed as bin volume fractions, *f*_bin1_, *f*_bin2_, and *f*_bin3_ (*D*_iso_<2.5 μm^2^/ms and *D*_Δ_^2^>0.25; *D*_iso_<2.5 μm^2^/ms and *D*_Δ_^2^<0.25; and *D*_iso_>2.5 μm^2^/ms and 0<*D*_Δ_^2^<1, respectively), which roughly represent WM, GM, and CSF signal fractions, respectively. Intra-voxel information from diffusion-relaxation correlations can be directly imaged by resolving the relaxation properties according to the above-defined bins (de Almeida Martins et al., 2020). We therefore included in this study the bin-resolved means of the *R*_1_ and *R*_2_ relaxation, e.g., E[*R*_1_]_bin1_, E[*R*_2_]_bin3_. The expectation values E[*x*] provide estimates of voxel averages of isotropic diffusivity *D*_iso_, squared diffusion anisotropy 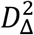, and relaxation rates *R*_1_ and *R*_2_, and are the closest analogues of mean diffusivity and fractional anisotropy in diffusion tensor imaging and monoexponential *R*_1_ and *R*_2_ relaxation rates in relaxation mapping. The frequency dependence parameters (Δ_ω⁄2*π*_[*x*]) describe how the diffusion parameters depend on the frequency of the diffusion encoding gradients. The bin fractions *f*_bin1_, *f*_bin2_, *f*_bin3_ are computed from the discrete *D*_iso_ and 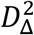 components and, thus, depend on their estimation accuracy. All of the above parameters are computed by taking the median over bootstrap replicas of the estimates.

### 2.5 Assessment of reliability and repeatability

#### 2.5.1 Intraclass correlation coefficient (ICC)

We used the intraclass correlation coefficient (ICC) as a measure of reliability. We used a single-measurement, absolute-agreement, two-way mixed-effects model (ICC(A, 1) in the convention of McGraw and Wong) for the computation (McGraw & Wong, 1996). MATLAB was used for the computation (MATLAB Release R2023a, The MathWorks, Inc., Natick, MA, USA). ICC estimates the proportion of variance due to differences between the participants compared to the total variance of the data. In other words, it is an estimate of the fraction of the total variance that is not due to measurement errors (or errors introduced by the processing pipeline, such as image registration inaccuracy). It characterizes whether differences between the participants can be detected from the noisy measurements. ICC can be improved by either decreasing the measurement error or by increasing the biological variability in the sample, such as by selecting participants with larger differences in age. The ICC ranges from 0 to 1, and while negative values are possible, they are treated as zeros. We use the classification used by Koo & Li (Koo & Li, 2016) to interpret the ICC values: poor reliability (ICC < 0.5), moderate reliability (0.5 ≤ ICC < 0.75), good reliability (0.75 ≤ ICC < 0.90), and excellent reliability (ICC ≥ 0.9).

#### 2.5.2 Within-subject coefficient of variation (CV_ws_)

We used the within-subject coefficient of variation (CV_ws_) as a measure of repeatability (Luque Laguna et al., 2020). It is the ratio of within-subject standard deviation *σ*_*ws*_ to the mean over subjects *μ*. We estimate CV_ws_ as

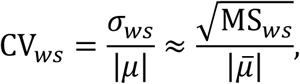

where MS_*ws*_ is the within-subject mean squares and 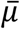 is the mean over participants and repetitions. Note that we take the absolute value of the mean because some of the MD-MRI parameters we use can take negative values. A within-subject coefficient of variation CV_*ws*_ *=* 0 indicates perfect repeatability.

### 2.6 Voxel-wise analysis

Discounting super-resolution and biophysical modeling-based approaches, voxel-wise analysis enables the most spatially specific analyses of MR images. For such analyses, MR images of the different subjects and time points are registered (transformed) to a common space, such as a study-specific template. Because the images are aligned, differences in MR signals between subjects (or time points) can be evaluated at each imaging voxel. However, this approach is sensitive to image registration inaccuracies and imaging noise. Accuracy of the image registration depends on the resolution, contrast, and signal-to-noise ratio of the images, as well as on the degree of anatomical differences between the subjects. The negative influences of image registration inaccuracies and measurement noise on voxel-wise analyses can be alleviated by using spatial filtering after image registration but this has a detrimental effect on the effective image resolution.

#### 2.6.1 Image registration

To enable the voxel-wise analysis, the MD-MRI parameter maps of the different participants at different time points were transformed to a common space (a study-specific template). Advanced Normalization Tools (Avants et al., 2011) (ANTs version 2.4.4, http://stnava.github.io/ANTs/, accessed on 14 June 2023) was used to compute and apply the image registrations. High-resolution (1-mm isotropic) T_1_-weighted MPRAGE images were used to create the study-specific template. First, repetition 1 and repetition 2 of each participant was registered to a mid-way space between the repetitions with an affine transform and averaged to avoid causing interpolation bias that would differ between each repetition (Reuter et al., 2012). Then, a study-specific template was formed from the test-retest average images of the different participants using a combination of affine and symmetric image normalization (SyN) transforms. The T_1_-weighted template image (1-mm isotropic resolution) was down-sampled to match the resolution of the MD-MRI images (2-mm isotropic resolution). Finally, the MD-MRI parameter maps of each participant and repetition were transformed to the 2-mm resolution template space by combining and applying the computed image transforms. Only one interpolation step was applied (after the application of all the image transforms in succession) to avoid the accumulation of image interpolation errors.

#### 2.6.2 Spatial filtering

After the transformation of the parameter maps to the template space, spatial filtering was applied to reduce the influence of inaccuracies caused by image registration. The choice of the optimal filtering function is nontrivial. We followed the approach of Caceres et al. to make that choice (Caceres et al., 2009). We applied Gaussian spatial filtering at different values of full width at half maximum (FWHM) for the kernel. When applying the filter, signal contributions from voxels outside of the brain mask were excluded. Then, we computed ICC for each voxel and the median of the ICC values of all brain voxels. A value of FWHM = 2 pixels (4 mm) was chosen for the voxel-wise analyses, as that value produced a substantial improvement in voxel-wise ICC values for all parameter maps without evidence of over-smoothing (Supplementary Fig. 1). Values of smoothing FWHM larger than 2 pixels decreased the median voxel-wise ICC values for some parameter maps, suggesting that the decrease in spatial specificity was outweighing the benefits associated with spatial smoothing.

#### 2.6.3 Brain-level statistics of voxel-wise metrics

After the application of spatial filtering (FWHM = 2 pixels = 4 mm), we computed reliability (ICC) and repeatability (CV_ws_) measures for each brain voxel. Violin plots were used to visualize the distributions of voxel-wise values of ICC and CV_ws_ over the whole brain. For some MD-MRI parameter maps, a small number of voxels had very high values of CV_ws_. The resulting distributions of voxel-wise values were very broad yet very sparse, making it difficult to produce plots that represented the correct shapes of the distributions. For those distributions of CV_ws_ that contained values higher than 100%, the MATLAB function “histcounts” (with automatically selected bins) was used to iteratively remove the voxels that were contained within histogram bins with a density of less than 0.1% of the total number of voxels. The resulting violin plots were labeled to indicate that they contained outliers that were removed prior to plotting. This outlier removal process was applied only for the violin plots of CV_ws_.

### 2.7 ROI-based analysis

In contrast to voxel-wise analysis, ROI-based analysis computes statistical measures within a chosen anatomical region. In our analysis, we compute the mean of each MD-MRI parameter map for each ROI. The resulting parameter estimates are thus spatially averaged based on anatomically delineated brain regions. Compared to voxel-wise analysis, ROI-based analysis is more robust as it is less susceptible to imaging noise and image registration inaccuracies. Voxel-wise analysis has gained popularity as an exploratory method, whereas ROI-based analysis is more common for (anatomically or functionally) hypothesis-driven studies.

#### 2.7.1 Whole brain segmentation

The spatially localized atlas network tiles (SLANT) method was used to perform whole brain segmentation (Huo et al., 2019) and obtain cortical and subcortical GM regions of interest (ROIs). Briefly, SLANT employs multiple independent 3D convolutional networks to segment the brain. Each of the networks is only responsible for a particular spatial region, thus the task of each network is simplified to focus on patches from a similar portion of the brain. Within this end-to-end pipeline, affine registration, N4 bias field correction, and intensity normalization are employed to roughly normalize each brain to the same space before segmentation. After each network performs its duty, the segmentation labels are fused together to form labels for 132 anatomical regions, primarily from subcortical and cortical areas, based on the BrainCOLOR protocol (https://mindboggle.info/braincolor/). The SLANT framework has shown high intra- and inter-scan protocol reproducibility (Xiong et al., 2019). The GM segmentation was performed to the T1-w MPRAGE images at 1-mm resolution. Finally, GM segmentations were down-sampled to 2-mm resolution to match the resolution of the MD-MRI scans.

The Johns Hopkins University DTI-based WM atlas (https://identifiers.org/neurovault.collection:264) was used to generate segmentations of white matter tracts. First, by using the participant-to-cohort average image transforms computed earlier (see Section 2.6.1), we formed a cohort-averaged 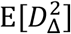 image (2-mm isotropic resolution). Then, we registered that image to the 2-mm resolution fractional anisotropy image of the WM atlas. Finally, we applied the inverses of the computed transforms (participant to cohort average and cohort average to WM atlas) to transform the ROIs of the WM atlas to the native space of each participant and repetition.

The segmented ROIs we focused on in our ROI analysis are shown in study-specific template space in Fig. 1. They covered different regions of the brain and included subcortical GM regions (basal ganglia and thalamus), cortical GM regions (middle occipital gyrus and inferior frontal gyrus), commissural tracts (genu, body, and splenium of the corpus callosum (CC)), projection tracts (internal capsule and corona radiata), and association tracts (sagittal stratum and external capsule). For each ROI that had a separate ROI for the left and right hemisphere of the brain, we combined the left and the right side to create a single ROI. To ascertain that each image voxel only belonged to one ROI, we removed from the GM ROIs those voxels that belonged to any of the WM ROIs. Finally, we removed all voxels with *f*_bin3_ > 0.2 (estimated CSF signal fraction > 0.2) from the ROIs. Some of the ROIs, shown for the different repetitions and participants in native space, are shown in Supplementary Fig. 2.

**Figure 1.**
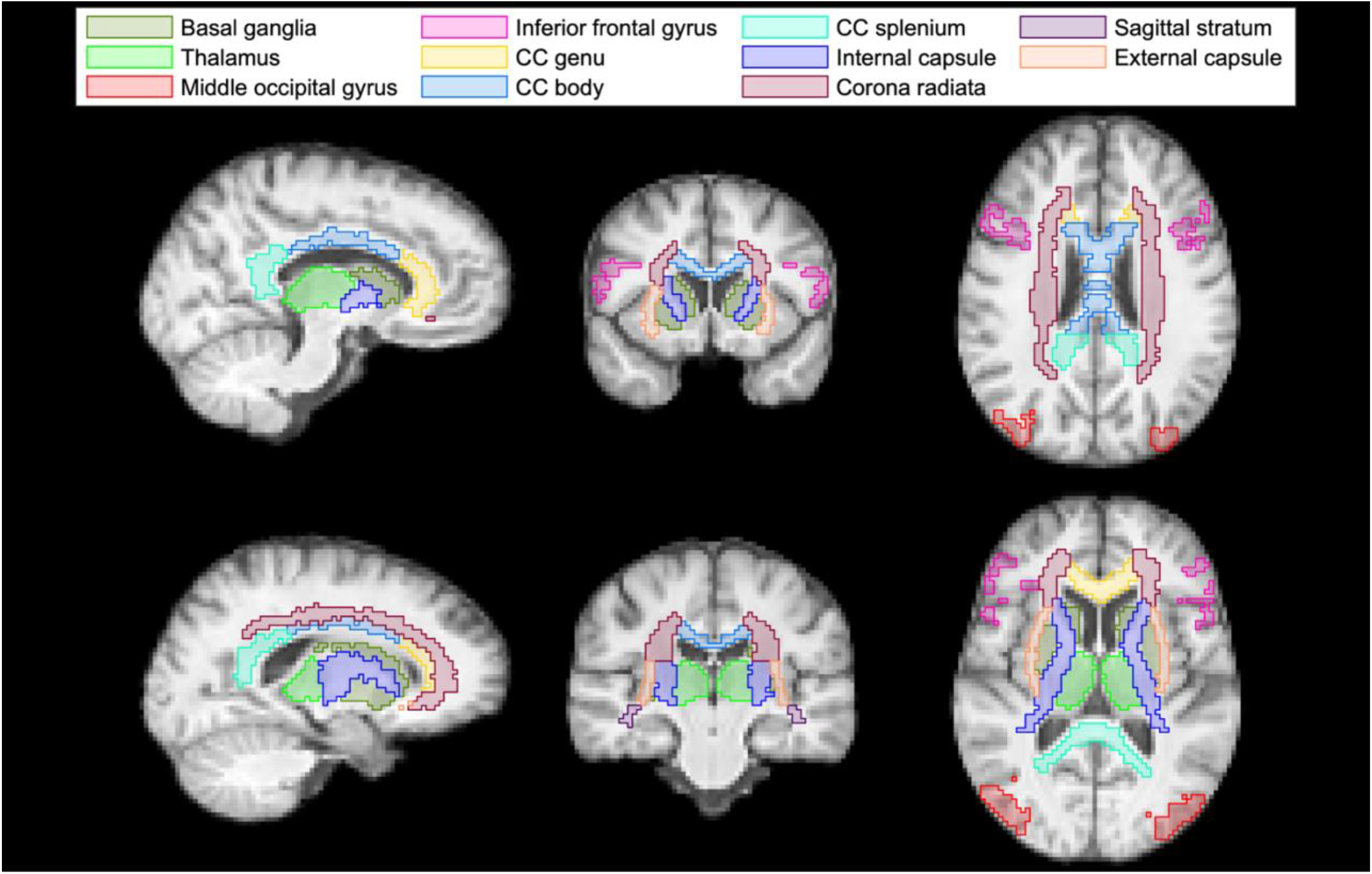
Regions of interest (ROIs) used in the ROI analysis as shown on the cohort-averaged T_1_-weighted template. Subcortical (basal ganglia and thalamus) and cortical (middle occipital gyrus and inferior frontal gyrus) gray matter ROIs were produced using the spatially localized atlas network tiles method (SLANT). Commissural (genu, body, and splenium of the corpus callosum (CC)), projection (internal capsule and corona radiata), and association (sagittal stratum and external capsule) white matter tracts were produced using the Johns Hopkins University diffusion tensor imaging-based white matter atlas.

#### 2.7.2 Estimation of ICC and CV_ws_

The estimation of ICC (reliability) and CV_ws_ (repeatability) values for each ROI and MD-MRI parameter map was performed in the native space of each participant and repetition to avoid unnecessary image interpolation. We employed an approach typical to ROI analysis, that is, we averaged the parameter value over the voxels within each ROI. Then, ICC and CV_ws_ were computed from those ROI-averaged parameter values. Thus, we get estimates for the reliability and repeatability of ROI-averaged MD-MRI parameter maps.

#### 2.7.3 Bland-Altman plots

Bland-Altman plots (Bland & Altman, 1986) were used to assess the presence of a bias between the test and retest measurement for each MD-MRI parameter map. In a Bland-Altman plot, the difference between the test and retest measurement is plotted as a function of the mean of the test and the retest measurement for each data point (in our case, each voxel). To achieve this, the parameter maps and ROIs of each participant and repetition were transformed to the cohort template space. In the cohort template space, a template ROI was formed by the majority vote of the ROIs of the participants and repetitions (at least 50% agreement between the participants and repetitions was needed for a voxel to belong to a certain ROI). Each voxel in the cohort template space that belonged to any of the ROIs included in our ROI analysis was included as a datapoint in the Bland-Altman plot, for each participant, resulting in 152620 data points (15262 voxels, 10 participants). The MATLAB class “densityScatterChart” (https://github.com/MATLAB-Graphics-and-App-Building/densityScatterChart, accessed July 17, 2023) was used to plot the density scatter plots. The mean of the test-retest difference over all datapoints (voxels and participants), along with the 95% confidence interval of the observations, were also computed and plotted for each MD-MRI parameter. Finally, we reported the relative mean test-retest difference as a percentage of the grand mean (mean over repetitions, participants, and voxels).

## 3 Results

### 3.1 Profiling of white matter, cortical, and subcortical regions in the human brain

Leveraging the rich information content of the voxelwise **D**(*ω*)-R_1_-R_2_ distributions, we first provide robust microstructural profiling across various brain regions, based on the average data from the 20 scans in our study. The ROIs used in this study are shown in Fig. 1, and projections from full **D**(*ω*)-R_1_-R_2_ distributions in six representative ROIs, comprising two cortical and two subcortical regions, along with two WM tracts are shown in Fig. 2.

**Figure 2.**
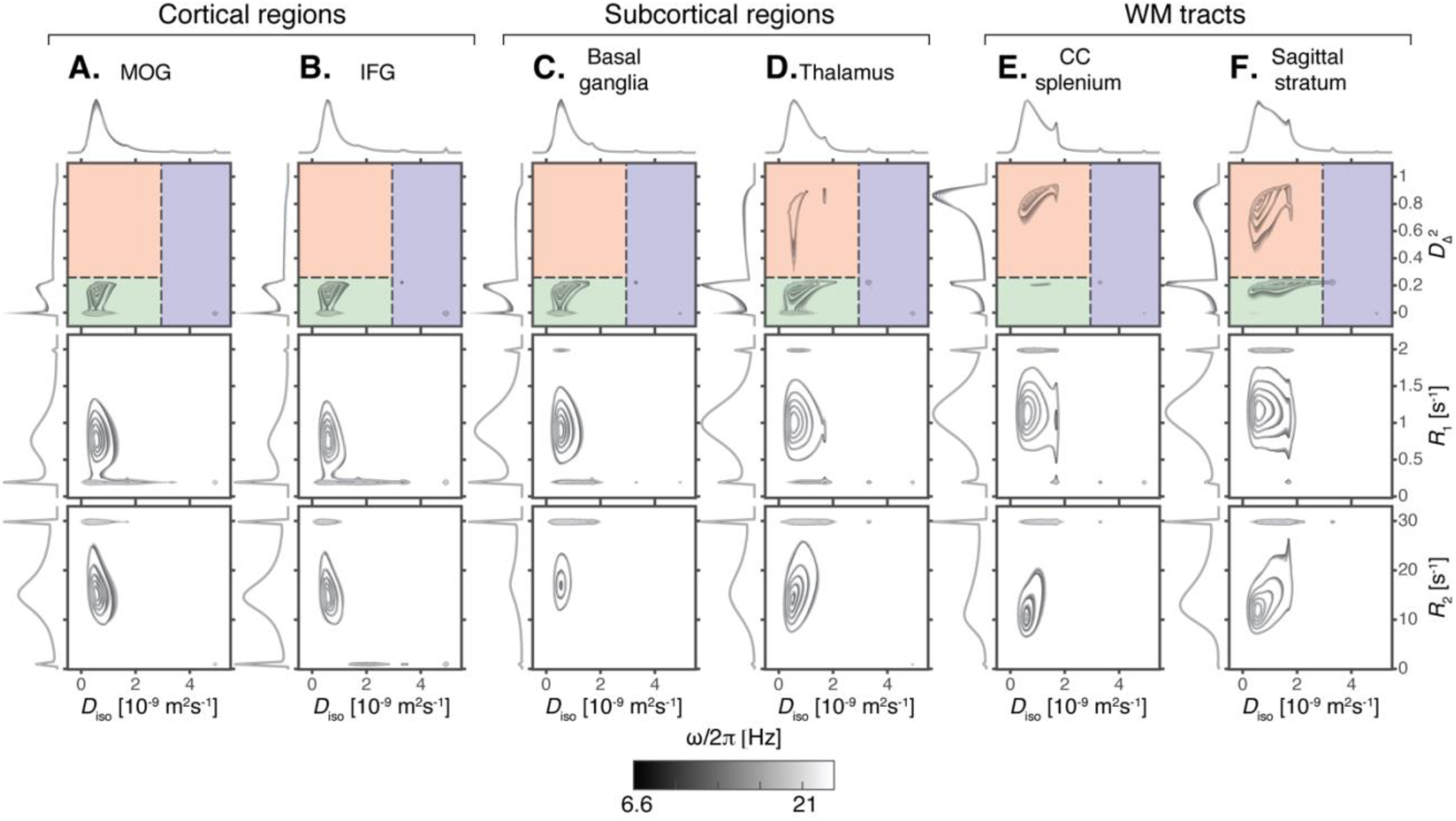
Projections of estimated **D**(*ω*)-R_1_-R_2_ distributions in different regions of interest (ROI). The displayed 2D projections include 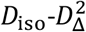 (top row), *D*_iso_-*R*_1_ (middle row), and *D*_iso_-*R*_2_ (bottom row) distributions. Different grayscale contour lines represent different centroid frequencies for the diffusion encoding gradients (*ω*⁄2*π =* 6.6 − 21 Hz). Colored areas in the 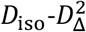 distribution plots represent bins that approximately correspond to white matter (red bin), gray matter (green bin), and cerebrospinal fluid (blue bin). Each distribution was averaged over the 10 participants, 2 repetitions, and the different voxels included in each ROI. (A) Middle occipital gyrus (MOG), (B) Inferior occipital gyrus (IFG), (C) Basal ganglia, (D) Thalamus, (E) Splenium of the corpus callosum (CC), (F) Sagittal stratum.

For each ROI, the distributions are visualized as projections onto the 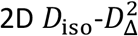, *D*_iso_-*R*_1_, and *D*_iso_-*R*_2_ planes for five frequencies ω/2*π* between 6.6 and 21 Hz. The well-known differences in tissue microstructure are most clearly manifested in the 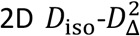 projection (Topgaard, 2019), which is therefore used to define three sub-voxel bins, as illustrated by the red, green, and blue overlays in Fig. 2. The bin signal fractions, *f*_bin1_, *f*_bin2_, and *f*_bin3_, evaluated at the lowest frequency (ω/2*π* = 6.6 Hz), are obtained by integrating over these spectral regions.

The distributions in Fig. 2 reflect the microstructural differences of the respective regions and their sub-voxel tissue components, revealing differences in composition between cortical and subcortical GM, and WM. Specifically for the 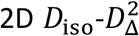 projections, cortical GM regions like the inferior frontal gyrus are mainly composed of morphologically isotropic tissue elements with low diffusivity (Fig. 2A-B, green shading). Subcortical regions like the thalamus present a wider distribution of tissue elements both in terms of their microscopic shape and size, with an additional highly anisotropic component (Fig. 2C-D, red shading); and WM tracts like the splenium of the corpus callosum are mainly composed of morphologically anisotropic and low diffusivity tissue elements (Fig. 2E-F).

The R_1_ and R_2_ relaxation rates can complement the microstructural picture with compositional information. The 2D *D*_iso_-*R*_1_ projections show a gradual increase in the R_1_ value of the main component from the cortical GM to the subcortical GM to the WM regions. A low *R*_1_ peak (∼0.2 1/s) was observed in cortical and subcortical regions, with a wider *D*_iso_ distribution for the former. A high R_1_ component can be seen, predominantly in WM, but also in subcortical regions. In the 2D *D*_iso_-*R*_2_ projections, distinct and more heterogeneous trends are observed compared to R_1_, with the highest R_2_ value of the main peak in the basal ganglia, followed by the thalamus and cortical regions, while WM tracts exhibit lower R_2_ values. A low *D*_iso_ with fast-relaxing R_2_ component was observed in all the ROIs, which could arise from noise and/or from fast-relaxing tissue components that are beyond the TE encoding range.

Diffusion frequency dependency is shown as grayscale contours on Fig. 2, and can be more readily observed from the 1D projections in each dimension. Although some frequency dependency can be seen for *D*_iso_, it is more pronounced when examining *D*_Δ_^2^. In all regions, frequency dependence of the microscopic anisotropy behaves in a similar manner, in which *D*_Δ_^2^ shifts towards lower values with increased frequency.

The voxel-wise multidimensional distributions can be used in different ways to extract information of interest. The most straightforward approach to achieve dimensionality reduction involves computing the per-voxel statistics, such as means E[x], for the different dimensions, assessed at one of the probed diffusion frequencies. This way, the means E[*D*_iso_], E[R_1_], and E[R_2_] correspond to conventional mean diffusivity (Basser et al., 1994), quantitative R_1_, and R_2_ (Weiskopf et al., 2021), respectively. The 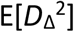 map is comparable to metrics used to quantify microscopic diffusion anisotropy (Lasič et al., 2014; Lawrenz et al., 2010; Shemesh et al., 2016). Expressing the diffusion frequency-dependence, rate of change with frequency (within the investigated range of 6.6–21 Hz) for the per-voxel means and variances can be used (Narvaez et al., 2022). Thus, in the Δ_*ω*⁄2*π*_ E[*D*_iso_] map, positive values indicate diffusion time dependency behavior suggestive of restriction (Aggarwal et al., 2012), while decreased anisotropy with higher frequency results in negative values in the 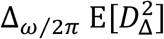 map. Per-voxel statistical values for the different dimensions, averaged within the ROIs in Fig. 2, are given in Table 1. The mean values agree with the expected microstructural differences between GM and WM regions, and also correspond with the observed trends of the 2D projections in Fig. 2.

**Table 1.** Mean (SD) values of multidimensional MRI parameters in select regions of interest.

| MD-MRI parameter | MOG | IFG | Basal ganglia | Thalamus | CC splenium | Sagittal stratum |
| --- | --- | --- | --- | --- | --- | --- |
| $E[D_{iso}]$ ($\mu\text{m}^2/\text{ms}$) | 0.95 (0.20) | 1.05 (0.22) | 0.93 (0.19) | 1.01 (0.17) | 1.09 (0.17) | 1.15 (0.14) |
| $E[D_{\Delta}^2]$ | 0.27 (0.11) | 0.249 (0.083) | 0.32 (0.12) | 0.38 (0.12) | 0.67 (0.15) | 0.522 (0.084) |
| $E[R_1]$ ($\text{s}^{-1}$) | 0.76 (0.13) | 0.71 (0.12) | 0.92 (0.12) | 0.99 (0.12) | 1.13 (0.14) | 1.15 (0.13) |
| $E[R_2]$ ($\text{s}^{-1}$) | 16.6 (2.3) | 15.2 (2.1) | 20.7 (3.7) | 17.9 (2.3) | 16.7 (2.3) | 16.6 (1.8) |
| $\Delta_{\omega/2\pi} E[D_{iso}]$ ($\mu\text{m}^2$) | 6.9 (5.9) | 0.8 (4.8) | 1.3 (4.7) | 1.0 (5.3) | 1.2 (3.3) | -0.2 (4.4) |
| $\Delta_{\omega/2\pi} E[D_{\Delta}^2]$ (ms) | -2.2 (1.2) | -2.0 (1.1) | -2.01 (0.99) | -2.5 (1.2) | -2.4 (1.6) | -2.7 (1.3) |
| $f_{bin1}$ | 0.36 (0.18) | 0.31 (0.15) | 0.41 (0.19) | 0.51 (0.18) | 0.84 (0.18) | 0.65 (0.13) |
| $f_{bin2}$ | 0.60 (0.16) | 0.62 (0.14) | 0.56 (0.20) | 0.46 (0.18) | 0.12 (0.16) | 0.30 (0.13) |
| $f_{bin3}$ | 0.027 (0.055) | 0.049 (0.068) | 0.010 (0.035) | 0.012 (0.039) | 0.015 (0.044) | 0.010 (0.034) |
| $E[R_1]_{bin1}$ ($\text{s}^{-1}$) | 0.63 (0.25) | 0.58 (0.23) | 0.82 (0.22) | 0.96 (0.17) | 1.15 (0.14) | 1.16 (0.14) |
| $E[R_1]_{bin2}$ ($\text{s}^{-1}$) | 0.78 (0.13) | 0.74 (0.12) | 0.88 (0.18) | 0.94 (0.21) | 0.41 (0.49) | 1.03 (0.29) |
| $E[R_1]_{bin3}$ ($\text{s}^{-1}$) | 0.06 (0.13) | 0.11 (0.16) | 0.020 (0.075) | 0.03 (0.12) | 0.04 (0.15) | 0.05 (0.17) |
| $E[R_2]_{bin1}$ ($\text{s}^{-1}$) | 16.3 (4.7) | 15.0 (4.4) | 20.6 (4.6) | 17.9 (2.6) | 16.1 (2.4) | 15.5 (2.0) |
| $E[R_2]_{bin2}$ ($\text{s}^{-1}$) | 15.1 (2.0) | 14.3 (1.9) | 18.4 (4.5) | 15 (4) | 5.7 (6.9) | 14.1 (4.1) |
| $E[R_2]_{bin3}$ ($\text{s}^{-1}$) | 0.8 (2.6) | 1.4 (3.0) | 0.5 (2.6) | 0.7 (2.9) | 0.8 (2.9) | 1.0 (3.5) |

An alternative way to extract information from the high dimensional voxel-wise data is to perform integration across a predefined parameter range, akin to a “spectral” ROI. The obtained integral value, ranging from 0 to 1, indicates the signal fraction within a particular multidimensional distribution. This voxel-wise computation enables the creation of an image reflecting the selected spectral component (Benjamini & Basser, 2020; Benjamini, Bouhrara, et al., 2021; de Almeida Martins et al., 2021; Martin et al., 2021). In this study, we used the previously suggested spectral ROIs shown in Fig. 2 (de Almeida Martins et al., 2020; Topgaard, 2019), to summarize the WM, GM, and CSF contributions. Per-voxel signal fractions, averaged within the ROIs in Fig. 2, are given in Table 1.

We can further leverage the derived sub-voxel information from the 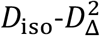 plane bins illustrated in Fig. 2 (i.e., *f*_bin1_, *f*_bin2_, and *f*_bin3_) and use it to resolve relaxation parameters based on their diffusion characteristics. Summarized values expressing diffusion-relaxation correlation via the (diffusion-) bin-resolved means of R_1_ and R_2_ relaxation properties are captured with the bin-resolved mean relaxation metrics, which can also be found in Table 1.

To complement the values in Table 1, means and standard deviations from all ROIs of each MD-MRI parameter, computed over all participants, repetitions, are shown in Supplementary Fig. 3.

### 3.2 Test–retest reliability and repeatability

We hypothesized that *in vivo* human brain **D**(*ω*)-*R*_1_-*R*_2_ MD-MRI parameter estimates acquired using a clinically achievable protocol are repeatable and reliable. Below, we evaluate this hypothesis for two prevalent image analysis approaches: voxel-wise and ROI-based. We employ the intraclass correlation coefficient (ICC) to assess the reliability, the within-subject coefficient of variation (CV_ws_) to quantify the repeatability, and Bland-Altman plots to estimate the bias of the MD-MRI parameter maps.

#### 3.2.1 Voxel-wise analysis

We transformed MD-MRI parameter maps to the cohort template space and estimated ICC (reliability) and CV_ws_ (repeatability) values for each brain voxel, after applying spatial filtering to the maps to reduce the effects of image misalignment. Parametric maps of mean values and signal fractions are shown in Fig. 3. Figure 3A shows a representative image slice of the MD-MRI parameter maps from a single subject. The maps are shown after transformation to the cohort template space but without the application of spatial filtering to give a representative example of the image quality. Fig. 3B shows the cohort averages (10 participants, 2 repetitions) for those MD-MRI parameter maps. Voxel-wise ICC and CV_ws_ values are shown in Figs. 3C and 3D, respectively. These were computed after applying a Gaussian spatial filter to the maps (FWHM = 2 pixels = 4 mm). Supplementary Fig. 4 shows ICC and CV_ws_ values computed with no spatial filtering applied to the parameter maps.

**Figure 3.**
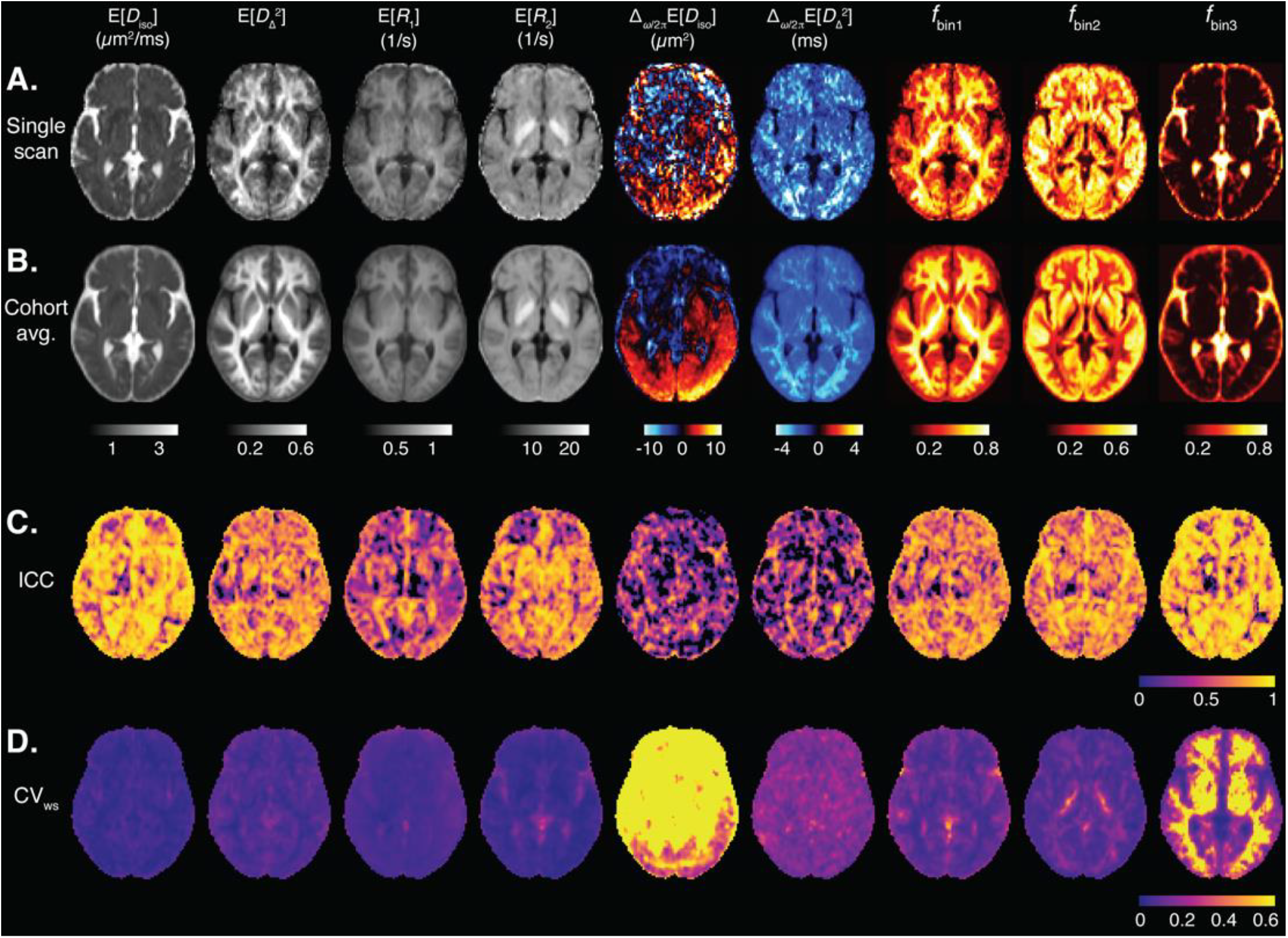
Voxel-wise MD-MRI parameter means E[x] and signal fraction maps, and their reliability and repeatability measures. Means E[x] and signal fraction maps, shown for (A) a representative single participant, and for a (B) cohort-average. Voxel-wise (C) intraclass correlation (ICC; reliability), and (D) within-subject coefficient of variation (CV_ws_; repeatability), computed for each parameter map. Spatial smoothing was applied to each map before the ICC and CV_ws_ computations. The ICC and CV_ws_ values demonstrate good reliability and repeatability for the first order statistical parameters E[*x*] of *D*_iso_, 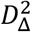, *R*_1_, and *R*_2_, and low reliability and repeatability for the dependence on diffusion gradient centroid frequency Δ_ω⁄2*π*_[*x*] of the diffusion parameters. The bin volume fractions *f*_bin1_ and *f*_bin2_ demonstrate high reliability and repeatability. The bin signal fraction *f*_bin3_ shows low repeatability in white matter but high reliability and repeatability elsewhere.

Maps of relaxation parameters resolved based on their diffusion characteristics are depicted in Fig. 4. Each map utilizes two orthogonal scales: the brightness intensity indicates the relative signal fraction, while the color scale denotes the specific relaxation property value. Voxel-wise ICC and CV_ws_ values are shown in Figs. 4C and 4D, respectively.

**Figure 4.**
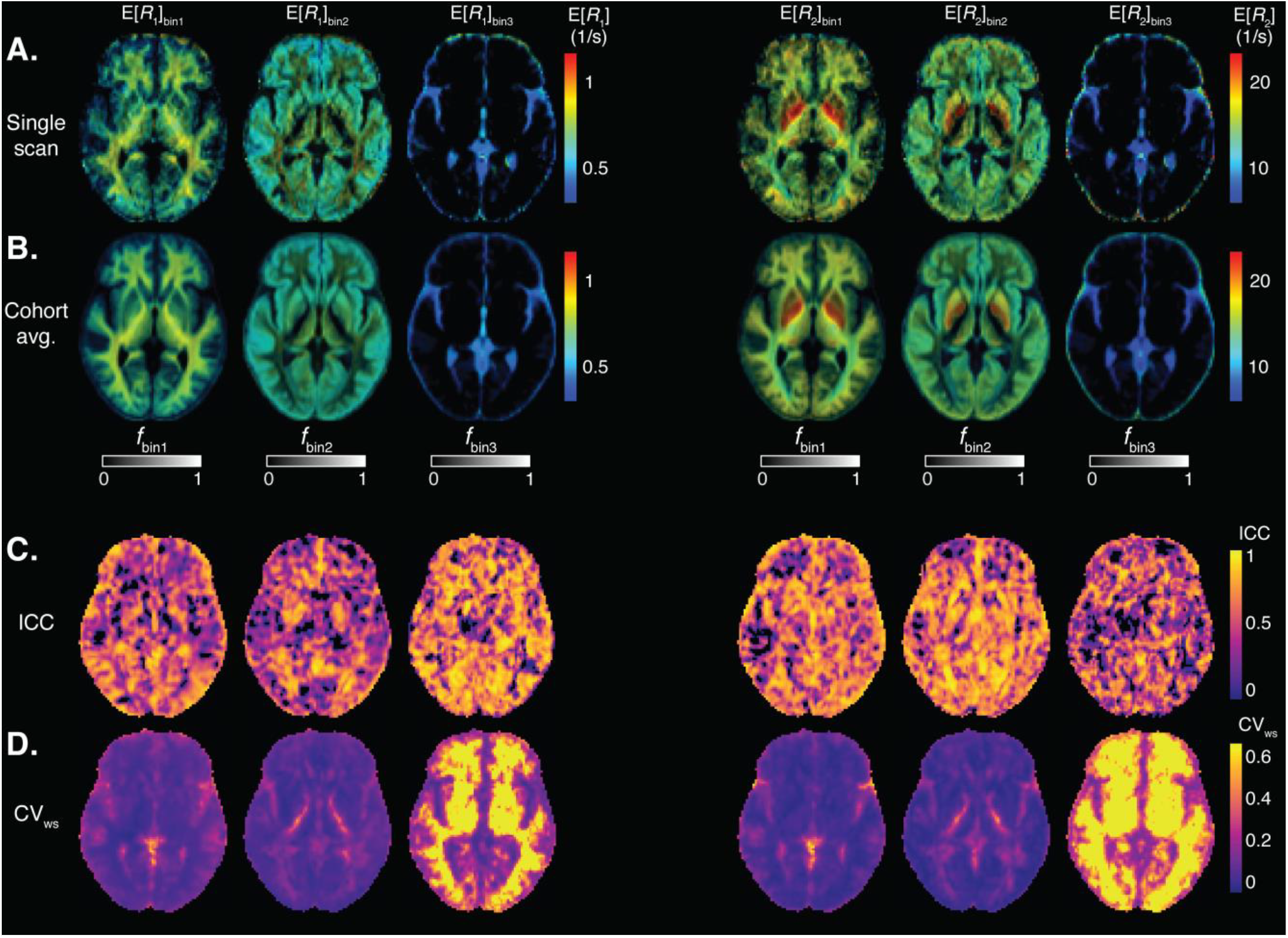
Voxel-wise MD-MRI parameter for bin-resolved relaxation values, and their reliability and repeatability measures. MD-MRI parameter R_1_ and R_2_ means, resolved according to their respective diffusion bin fractions, shown for (A) a representative single participant, and for (B) a cohort-average. Voxel-wise (C) intraclass correlation coefficient (ICC; reliability), and (D) within-subject coefficient of variation (CV_ws_; repeatability), computed for each parameter map. Spatial smoothing was applied to each map before the ICC and CV_ws_ computations. The ICC and CV_ws_ values demonstrate good reliability and repeatability except for *f*_bin3_, which shows low repeatability in white matter but high reliability and repeatability elsewhere.

Table 2 shows median and interquartile range values of voxel-wise ICC and CV_ws_ for each MD-MRI parameter map. With most MD-MRI parameter maps, high ICC values (high reliability) corresponded to low CV_ws_ values (high repeatability), as could be expected. E[*D*_iso_], 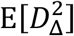, and E[*R*_2_] exhibited both high ICC values as well as low CV_ws_ values. E[*R*_1_] showed one of the lowest CV_ws_ values but also low ICC values, suggesting low relative measurement error but also small biological variability between the participants. The estimated signal fraction parameter *f*_bin3_ had relatively high ICC but high CV_ws_ values as well. The high CV_ws_ values of *f*_bin3_ were concentrated in WM regions. In those regions, *f*_bin3_ values are close to zero, and so even minute differences in *f*_bin3_ between repetitions lead to high CV_ws_ values. We also estimated the distributions of the ICC and CV_ws_ values over all brain voxels. Supplementary Fig. 5 shows the distributions of ICC and CV_ws_ values in the brain with different levels of spatial filtering applied to the parameter maps.

**Table 2.**
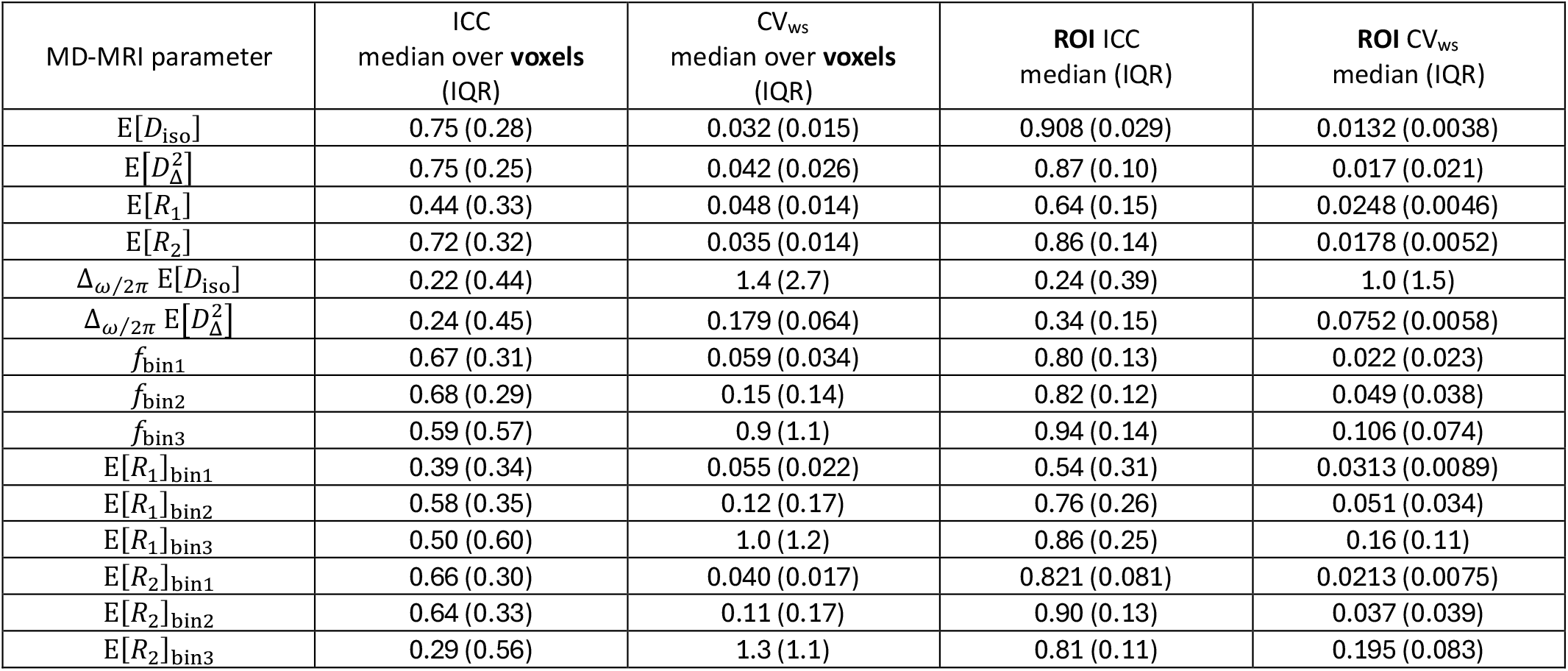
Summary of voxel-wise and region of interest (ROI)-based reliability (intraclass correlation coefficient, ICC) and repeatability (within-subject coefficient of variation, CV_ws_) results. For voxel-wise results, median and interquartile range (IQR) of ICC and CV_ws_ values were taken over all brain voxels. For ROI-based results, ICC and CV_ws_ were computed for each ROI-averaged multidimensional (MD)-MRI parameter map, and the median and IQR values were computed over the ROI-wise ICC and CV_ws_ values.

| MD-MRI parameter | ICC<br>median over **voxels**<br>(IQR) | $CV_{ws}$<br>median over **voxels**<br>(IQR) | **ROI** ICC<br>median (IQR) | **ROI** $CV_{ws}$<br>median (IQR) |
| --- | --- | --- | --- | --- |
| $E[D_{iso}]$ | 0.75 (0.28) | 0.032 (0.015) | 0.908 (0.029) | 0.0132 (0.0038) |
| $E[D_{\Delta}^2]$ | 0.75 (0.25) | 0.042 (0.026) | 0.87 (0.10) | 0.017 (0.021) |
| $E[R_1]$ | 0.44 (0.33) | 0.048 (0.014) | 0.64 (0.15) | 0.0248 (0.0046) |
| $E[R_2]$ | 0.72 (0.32) | 0.035 (0.014) | 0.86 (0.14) | 0.0178 (0.0052) |
| $\Delta_{\omega/2\pi} E[D_{iso}]$ | 0.22 (0.44) | 1.4 (2.7) | 0.24 (0.39) | 1.0 (1.5) |
| $\Delta_{\omega/2\pi} E[D_{\Delta}^2]$ | 0.24 (0.45) | 0.179 (0.064) | 0.34 (0.15) | 0.0752 (0.0058) |
| $f_{bin1}$ | 0.67 (0.31) | 0.059 (0.034) | 0.80 (0.13) | 0.022 (0.023) |
| $f_{bin2}$ | 0.68 (0.29) | 0.15 (0.14) | 0.82 (0.12) | 0.049 (0.038) |
| $f_{bin3}$ | 0.59 (0.57) | 0.9 (1.1) | 0.94 (0.14) | 0.106 (0.074) |
| $E[R_1]_{bin1}$ | 0.39 (0.34) | 0.055 (0.022) | 0.54 (0.31) | 0.0313 (0.0089) |
| $E[R_1]_{bin2}$ | 0.58 (0.35) | 0.12 (0.17) | 0.76 (0.26) | 0.051 (0.034) |
| $E[R_1]_{bin3}$ | 0.50 (0.60) | 1.0 (1.2) | 0.86 (0.25) | 0.16 (0.11) |
| $E[R_2]_{bin1}$ | 0.66 (0.30) | 0.040 (0.017) | 0.821 (0.081) | 0.0213 (0.0075) |
| $E[R_2]_{bin2}$ | 0.64 (0.33) | 0.11 (0.17) | 0.90 (0.13) | 0.037 (0.039) |
| $E[R_2]_{bin3}$ | 0.29 (0.56) | 1.3 (1.1) | 0.81 (0.11) | 0.195 (0.083) |

#### 3.2.2 ROI analysis

The ROIs used in the analysis are shown in Fig. 1. Within each ROI and for each participant and repetition, the mean MD-MRI parameter map was computed, from which ICC (reliability) and CV_ws_ (repeatability) were estimated. As the parameters are averaged over the ROIs, the number of voxels in each ROI affects the quality of the metrics. All ROIs in this study had comparable volume (∼1500 voxels), with the exception of the corona radiata (∼3500 voxels) and the sagittal stratum (∼500 voxels). The number of voxels for each participant and repetition in each ROI is shown in Supplementary Fig. 6.

Reliability and repeatability values for different MD-MRI parameter maps and ROIs are plotted in Fig. 5, and the individual values given in Supplementary Fig. 7. Median and interquartile range over all ROIs for the ROI-based ICC and CV_ws_ values are shown in Table 2. These findings indicate, as expected, that ROI-averaged MD-MRI parameters are more reliable and repeatable than their voxel-wise counterparts. The relationship between high reliability (high ICC) and high repeatability (low CV_ws_) in the ROI-based analysis was similar to the results of the voxel-wise analysis, except in the case of *f*_bin3_.

**Figure 5.**
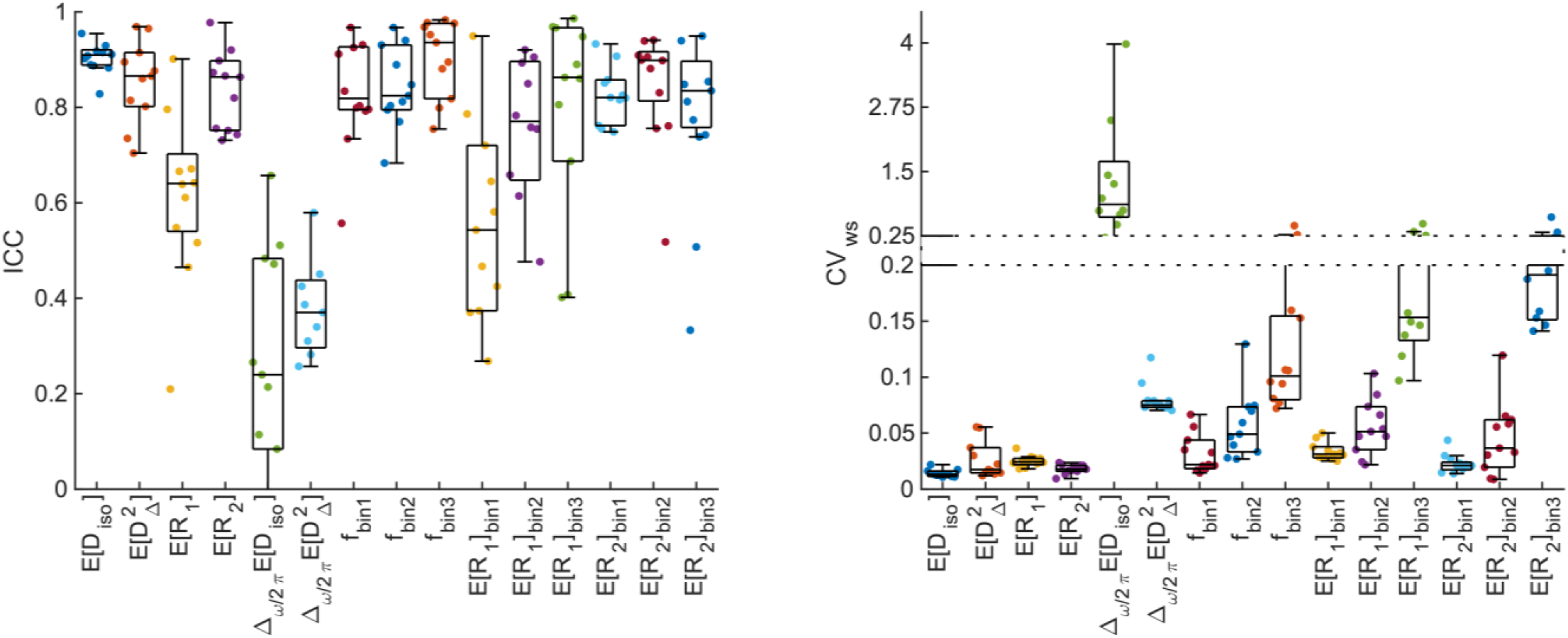
Reliability and repeatability of region-of-interest (ROI)-averaged MD-MRI parameter maps. (A) Intraclass correlation coefficient (ICC; reliability). (B) Within-subject coefficient of variation (CV_ws_; repeatability). Generally, ROI-based ICC values were higher and CV_ws_ values lower than those of individual voxels (Figures 3 and 4; Table 2). That was expected due to the increase in signal-to-noise ratio achieved through averaging. Numeric values for each ROI are given in Supplementary Fig. 7.

The reliability (ICC) of *f*_bin3_ benefited more from the ROI-averaging than the other MD-MRI parameters, achieving the highest median ROI-based reliability (ICC = 0.94) out of all the MD-MRI parameters even though its median repeatability was the 4^th^ worst (CV_ws_ = 0.106). As in the voxel-wise analysis, the ROI-averaged values of E[*D*_iso_], 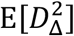, and E[*R*_2_] displayed both high reliability and high repeatability.

We assessed the presence of bias between the test and the retest measurement using Bland-Altman plots comprising the different participants and the voxels included in the analyzed ROIs (Fig. 6). For each of the MD-MRI parameters, the 95% confidence interval for the observations included the zero-bias level. The relative mean differences between the test and the retest measurement for the different parameters were E[*D*_iso_]: 0.55%, 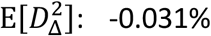, E[*R*_1_]: -0.66%, E[*R*_2_]: -0.65%, Δ_ω⁄2*π*_E[*D*_iso_]: 11%, 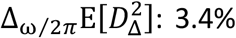, *f*_bin1_: -0.60%, *f*_bin2_: 1.1%, *f*_bin3_: -1.7%, E[*R*_1_]_bin1_: -0.84%, E[*R*_1_]_bin2_: -0.51%, E[*R*_1_]_bin3_: 2.0%, E[*R*_2_]_bin1_: -0.96%, E[*R*_2_]_bin2_: 0.51%, E[*R*_2_]_bin3_: 6.0%. Those biases were, on average, about one order of magnitude smaller (range: 4 to 140 times smaller) than the median voxel-wise within-subject coefficients of variation CV_ws_ (Table 2).

**Figure 6.**
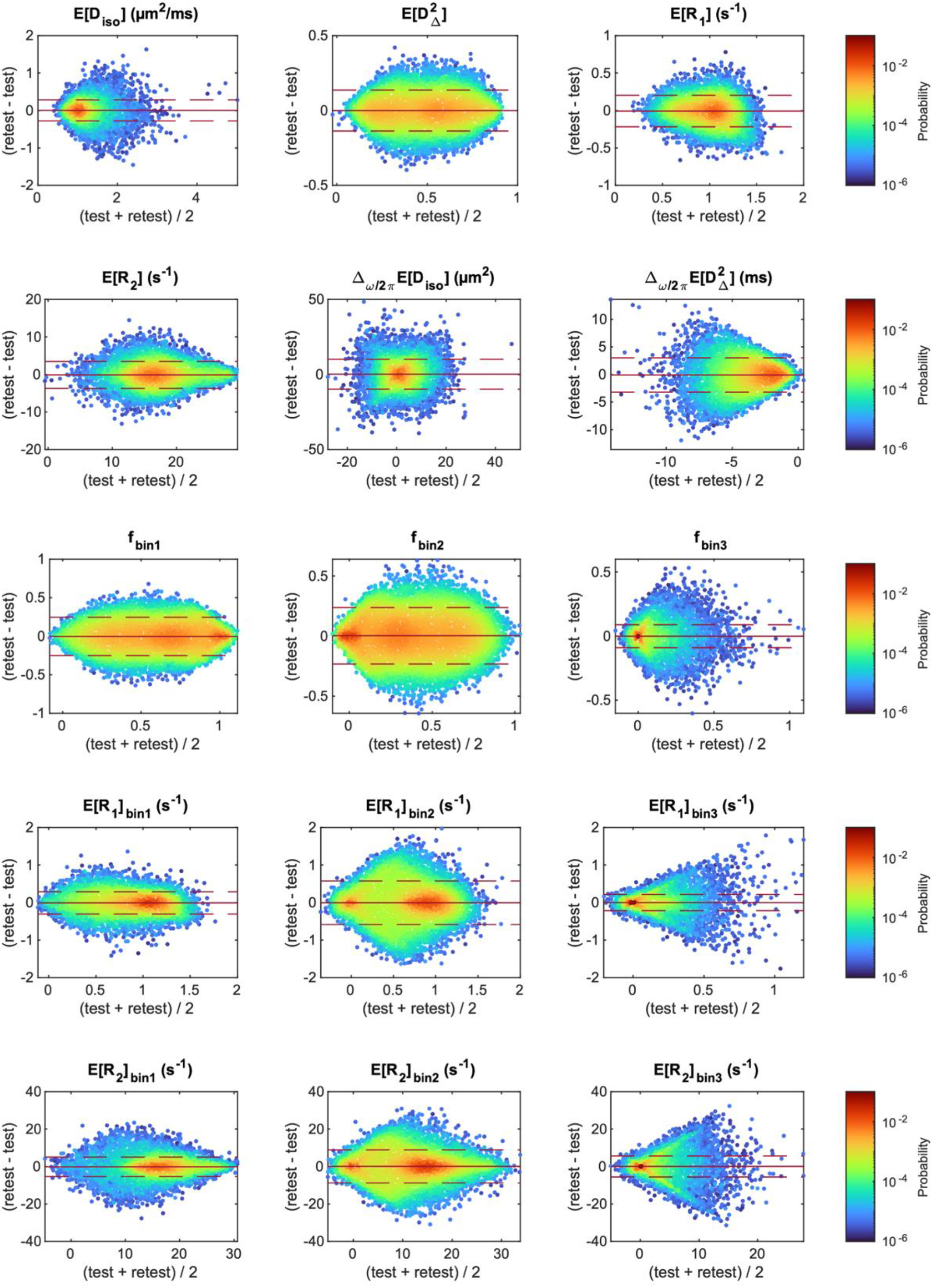
Bland-Altman density plots for the different MD-MRI parameter maps. In each plot, all the voxels that were included in the region of interest analysis are included as data points, for all participants. The mean (solid red line) and the 95% confidence interval of the observations (1.96 × standard deviation; dashed red line) are shown for each plot.

## 4 Discussion

Diffusion-relaxation correlation MD-MRI is a method that allows to acquire rich information about tissue microstructure by exploring the correlations between different MRI contrasts. This framework replaces voxel-averaged quantities with multidimensional distributions of those quantities that allows to selectively extract and map specific diffusion-relaxation spectral ranges. As a result, it has the potential of achieving high sensitivity and specificity towards detecting subtle changes that would have been otherwise averaged-out. The purpose of this work was to methodologically characterize different brain regions in terms of their multicomponent diffusion-relaxation properties, interpret differences and similarities between them, and to assess the framework’s overall repeatability. We compared WM tracts, cortical GM, and subcortical GM regions, and found pronounced differences in their diffusion-relaxation profile, reflecting distinct differences in their microscopic morphological content. We then applied a test-retest paradigm to estimate the repeatability and reliability of parameters derived from the 40-min whole brain *in vivo* MD-MRI framework, and showed them to be comparable with known DTI reproducibility.

Results from comprehensive profiling of representative WM, cortical GM, and subcortical GM regions are shown in Fig. 2 and Table 1. Given the extensive history of DTI studies, it is not surprising that the MD-MRI data allowed to easily differentiate between WM and GM regions based on their different diffusion anisotropy distributions. However, we demonstrated here that meaningful differences can be gleaned even between WM tracts: compared with the sagittal stratum, the CC splenium had a lower content of microscopically isotropic and restricted components (i.e., *f*_bin2_, See Table 1), and had, in addition, a narrower R_2_ distribution. These differences may arise from the layered structure of the sagittal stratum (Di Carlo et al., 2019) that makes it more microstructurally heterogeneous compared with the CC.

Our findings in GM revealed distinct microstructural attributes that differentiate between subcortical and cortical regions. Compared with the cortical regions, the thalamus and basal ganglia contained a higher content of microscopically anisotropic components (i.e., *f*_bin1_, see Table 1), and a wider distribution of diffusivities, skewed towards larger values. These differences in microscopic anisotropy and in diffusivity are likely to be driven, respectively, by the presence of fiber tracts and the relatively large subcortical nuclei (Morel, 2007). In addition to the diffusion properties, cortical and subcortical GM regions showed differences in their *R*_1_ distributions.

While we were able to apply the MD-MRI framework to perform a microstructural study of the human brain thus demonstrating its potential, there are two factors that may introduce unwanted variability to its outcomes: the acquisition parameter space is sparsely sampled to keep the scan time feasible, and the parameter estimation is an ill-posed inverse problem. Until now, there has been no evaluation of MD-MRI parameter reproducibility in the living human brain. Therefore, we investigated the reliability (ICC) and repeatability (CV_ws_) of estimated MD-MRI parameters using a test-retest paradigm. The CV_ws_ quantifies the relative error between repetitions of the same measurement (Luque Laguna et al., 2020). Thus, high repeatability means that the difference between the repetitions is small compared to the measured value. The ICC quantifies the fraction of variance that is not explained by errors, i.e., that is due to differences between the participants (Matheson, 2019). High reliability means that biological differences between participants can be detected reliably. Like CV_ws_, ICC depends on errors but, unlike CV_ws_, also on differences between participants. Thus, our results for reliability are representative of our sample of participants (ten healthy participants with a mean age of 48 years, standard deviation 14.4 years; 4 women). The results could be different for a different age or sex distribution or in the presence of disease or pathology.

We investigated the reliability and repeatability of both voxel-based and ROI-averaged MD-MRI parameter estimates. The MD-MRI distribution mean parameters all had median voxel-wise CV_ws_ values smaller than 5%, exhibiting good repeatability. Voxel-wise E[*D*_iso_] and 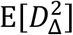, demonstrated good reliability (ICC = 0.75) and E[*R*_2_] moderate reliability (ICC = 0.72). However, E[*R*_1_] had poor reliability (ICC<0.50), which may be due to the relatively narrow range of R_1_ encoding in the acquisition protocol, leading to limited sensitivity. The ROI-averaged distribution means E[*D*_iso_], 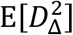, and E[*R*_2_] had high repeatability (CV_ws_ < 2%) and reliability (ICC > 0.85).

The diffusion gradient frequency dependency of mean isotropic diffusivity, Δ_*ω*⁄2*π*_ E[*D*_*iso*_], exhibited poor voxel-wise and ROI-averaged reliability and repeatability. 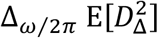 had poor reliability but reasonably good repeatability (voxel-wise CV_ws_ = 18%, ROI CV_ws_ = 7.5%). We believe that these results have two main reasons. First, diffusion gradient hardware limitations constrain the diffusion frequency range to centroid frequencies of 6.6–21 Hz, with the highest frequency achieved only at a low b-value of 0.5 ms/μm^2^, and thus restrict the ability to reliably decompose and estimate the expected time scales in the system. Using this frequency content range is expected to be particularly sensitive to the length scale of approximately 13 μm (Stepišnik, 1993; Woessner, 1963). While this length scale may be sufficiently prevalent in some brain regions, typical length scales in most of the human brain are significantly smaller. This mismatch between the sensitivity range of the MD-MRI acquisition and the underlying microstructure is a potential driver of the poor reproducibility we found. We believe that the second contributor to these findings is the way in which the frequency-dependence parameters are defined, i.e., as a subtraction of two parameters. Thus, they suffer from error propagation as inaccuracies are summated during subtraction. Furthermore, the absolute differences between the highest and lowest frequencies are very small due to the limited frequency range, leading to small numbers around 0, with reduced sensitivity. Based on these results, finding an alternative way to quantify frequency-dependence from the **D**(*ω*)-R_1_-R_2_ distributions is encouraged.

One of the unique features of the MD-MRI framework is the ability to access and visualize intravoxel information within the human brain by selectively integrating regions of the voxel-wise **D**(*ω*)-R_1_-R_2_ distributions. Consistent with previous studies, we delineated three *spectral* ROIs in the 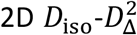 distribution space, roughly corresponding to WM, GM, and CSF, denoted as *bin1, bin2*, and *bin3*, respectively, as illustrated in Fig. 2. Voxel-wise application of partial integration based on these bins yields corresponding signal fraction maps, labeled as *f*_bin1_, *f*_bin2_, and *f*_bin3_. Voxel-wise reliability and repeatability analyses resulted in whole brain median ICC values of 0.67, 0.68, and 0.59, and CV_ws_ values of 6%, 15%, and 90%, for *f*_bin1_, *f*_bin2_, and *f*_bin3_, respectively. The ROI-based performance was better, with median ICC values ranging from 0.80 to 0.94, and CV_ws_ values in the range of 2-11%. We hypothesize that the relatively poor reproducibility of *f*_bin3_ was due to the small amounts of free water in non-CSF regions in the brain, leading to high relative errors in the estimated signal fractions. This effect is particularly exacerbated when reporting median values across the whole brain, with only a small portion of the voxels residing in CSF-containing regions, as can be seen in the almost binary-looking *f*_bin3_ map in Fig. 3. The overall good reproducibility of the intravoxel fraction maps is encouraging, especially because such sub-voxel component maps have demonstrated specificity towards pathology and have been used to characterize subtle processes noninvasively (Benjamini, Iacono, et al., 2021; Benjamini et al., 2022; Slator et al., 2019).

The sub-voxel spectral information can be further utilized to resolve relaxation parameters based on their diffusion characteristics, such as the bin-resolved mean relaxation rates E[*R*_1_]_bin1_, E[*R*_1_]_bin2_, E[*R*_1_]_bin3_, E[*R*_2_]_bin1_, E[*R*_2_]_bin2_, and E[*R*_2_]_bin3_, thus directly visualizing diffusion-relaxation correlation. Of those bin-resolved parameters, E[*R*_1_]_bin2_, E[*R*_1_]_bin3_, E[*R*_2_]_bin1_, and E[*R*_2_]_bin2_ had moderate median voxel-wise reliabilities (ICC > 0.5) and good ROI-averaged reliabilities (ICC > 0.75). Their repeatability values were relatively good (voxel-wise CV_ws_ 4-12%, ROI CV_ws_ 2-5%), except those of E[*R*_1_]_bin3_, which were poorer (voxel-wise CV_ws_ = 100%, ROI CV_ws_ = 16%). E[*R*_1_]_bin1_ had low voxel-wise reliability (ICC = 0.39), moderate ROI reliability (ICC = 0.54), and good repeatability (voxel-wise CV_ws_ = 5.5%, ROI CV_ws_ = 3.1%). Interestingly, E[*R*_2_]_bin3_ had poor voxel-wise reliability (ICC = 0.29) and repeatability (CV_ws_ = 130%) but good ROI-averaged reliability (ICC = 0.81). Taken together, these results suggest that MD-MRI scalar parameters that combine information from different contrasts (diffusion and relaxation) can be reliable and repeatable. However, parameters related to bin3 (CSF) provide a special case to consider. Such parameters had considerably lower, even very poor, median voxel-wise reliability and repeatability values. Averaging those parameters over an ROI greatly increased their reliability and repeatability values. The voxel-wise estimates could have been greatly influenced by the low amount of CSF in most tissues and by inaccuracies in image registration.

Comparing our reliability and repeatability results to previous studies would provide important context. While the current study is the first to report on reproducibility of MD-MRI, we can draw parallels with previous diffusion MRI variability investigations. Specifically, MD and FA in our study align with E[*D*_iso_] and 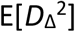, respectively. Most of the recent DTI studies investigated ICC and CV_ws_ in WM regions. Repeatability and reliability of DTI metrics in the CC reported, for MD and FA respectively, CV_ws_ ranges of 1-2.5% and 2.5-3%, and ICC ranges of 0.54-0.73 and 0.76-0.98 (Coelho et al., 2022; Fan et al., 2021; Grech-Sollars et al., 2015; Luque Laguna et al., 2020). Our investigation in the CC revealed, for E[*D*_iso_] and 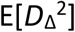 both an average CV_ws_ of 1%, and average ICC of 0.88 and 0.91, respectively. In addition to E[*D*_iso_] and 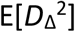, the reproducibility of E[R_1_] and E[R_2_] can also be compared with relaxometry studies in the CC, in which the CV_ws_ was established at under 1% for both T_1_ and T_2_ (Hagiwara et al., 2019), comparable with our reported values around 2%. Another rough comparison could be drawn between the free water fraction from the so-called ‘Standard Model’ (Novikov et al., 2019) and MD-MRI derived f_bin3_. A recent reproducibility study found CV_ws_ values of 9% and 15% in the CC and internal capsule, respectively (Coelho et al., 2022), comparable with CV_ws_ average values of 10% and 16% we reported here for f_bin3_. Notably, the reproducibility of MD-MRI diffusion measures in the subcortical regions we investigated surpassed those of reported DTI parameters. For instance, reliability for MD and FA in the basal ganglia (CV_ws_ of 4% and 10%) and in the thalamus (CV_ws_ of 10% and 8%) (Grech-Sollars et al., 2015) should be compared with the corresponding values of 2% for E[*D*_iso_] and 4% for E[*D*_Δ_ ^2^] reported in our study.

Our study has several limitations and assumptions. The MD-MRI signal representation used provides nonparametric distributions of diffusion and relaxation components and is based on the Gaussian phase distribution (GPD) approximation (Neuman, 1974; Stepišnik, 1981). Non-Gaussian diffusion effects that violate the GPD assumption (Jespersen et al., 2019) and induce microscopic kurtosis are possible (Novello et al., 2022). In this case, the microscopic kurtosis is not accounted for in our model, and its effect would split between variances of *D*_iso_ and *D*_Δ_^2^. Considering relaxation, quantitative comparisons of R_1_ and R_2_ across studies are hindered by fundamental considerations in pulse sequence and acquisition design. The recorded signal results from intricate interactions among partial excitation, relaxation processes, and proton pool exchange, each possessing distinct MR properties (Manning et al., 2021). These factors make R_1_ and, to a lesser extent, R_2_, dependent on specific pulse sequence parameters, slice thickness, and radiofrequency bandwidth. Furthermore, hardware limitations restricted the minimal echo time to 40 ms in this study, which may fully attenuate myelin water (R_2_ ∼ 100 s^-1^) (Manning et al., 2021). Future improvements in diffusion gradient frequency range, b-values, and echo time minimization could be achieved by utilizing next-generation diffusion gradients (Dai et al., 2023; Huang et al., 2021). When assessing the reproducibility results, we should keep in mind that the study included ten participants, and the results related to reliability are representative of that sample. Although consistent with the median sample size of six reported in technical MRI studies (Hanspach et al., 2021), a larger sample would likely yield more robust MD-MRI reproducibility values. Additionally, our focus on intrascanner reproducibility highlights the need for future investigations into inter-scanner assessment. Furthermore, while informative, voxel-wise reproducibility analysis requires image registration to a common space, introducing inaccuracies due to biological variation and image interpolation. Although we applied Gaussian spatial filtering to mitigate these effects (Caceres et al., 2009), complete elimination cannot be achieved. Similarly, the ROI-based analysis is affected by the accuracy of the segmentation results (Supplementary Fig. 2).

## 5 Conclusion

We established reliable quantification of relaxation and diffusion properties, including intravoxel information, using a clinically feasible, 40-min *in vivo* diffusion-relaxation correlation (**D**(*ω*)-R_1_-R_2_) multidimensional MRI scan in the human brain. This enabled us to methodologically explore brain regions in terms of their multicomponent properties and interpret differences and similarities between them. We were able to observe subtle variations between subcortical and cortical regions, demonstrating the microstructural sensitivity of the MD-MRI framework. The possibilities of using the rich spectral information in the MD-MRI data to create biomarkers that show high sensitivity to specific cell types or pathologies should be further investigated. To ensure the reproducibility and adoption of such biomarkers, future studies should also investigate the inter-scanner reproducibility and the harmonization of MD-MRI datasets.

## Data and Code Availability

The data that support the findings of this study are available on request from the corresponding author. The data are not publicly available due to privacy or ethical restrictions. Code to process the MD-MRI data is freely available as implemented in the multidimensional diffusion MRI toolbox (https://github.com/markus-nilsson/md-dmri).

## Author Contributions

Eppu Manninen: Conceptualization, Methodology, Software, Formal Analysis, Writing – Original Draft, Visualization; Shunxing Bao: Software, Resources, Writing - Review & Editing; Bennett A. Landman: Software, Resources, Writing - Review & Editing; Yihong Yang: Resources, Funding acquisition, Writing Review & Editing; Daniel Topgaard: Conceptualization, Methodology, Software, Writing - Review & Editing; Dan Benjamini: Conceptualization, Methodology, Software, Investigation, Resources, Writing Original Draft, Visualization, Supervision, Project administration, Funding acquisition.

## Funding

This work was funded by the Intramural Research Programs of the National Institute on Aging (NIA) and the National Institute on Drug Abuse (NIDA) of the National Institutes of Health (NIH).

## Declaration of Competing Interest

The authors declare no conflict of interest.

## Acknowledgements

The authors would like to thank Mr. Phil Cholak for facilitating the MRI scans.

## Notes

### Competing Interest Statement

The authors have declared no competing interest.

## References

Aggarwal, M., Jones, M. V., Calabresi, P. A., Mori, S., & Zhang, J. (2012). Probing mouse brain microstructure using oscillating gradient diffusion MRI. Magnetic Resonance in Medicine, 67(1), 98–109. 10.1002/mrm.22981

Assaf, Y., & Basser, P. J. (2005). Composite hindered and restricted model of diffusion (CHARMED) MR imaging of the human brain. NeuroImage, 27(1), 48–58. 10.1016/j.neuroimage.2005.03.042

Avants, B. B., Tustison, N. J., Song, G., Cook, P. A., Klein, A., & Gee, J. C. (2011). A reproducible evaluation of ANTs similarity metric performance in brain image registration. NeuroImage, 54(3), 2033–2044. 10.1016/j.neuroimage.2010.09.025

Basser, P. J., Mattiello, J., & LeBihan, D. (1994). MR diffusion tensor spectroscopy and imaging. Biophysical Journal, 66(1), 259–267. 10.1016/S0006-3495(94)80775-1

Benjamini, D. (2020). Nonparametric Inversion of Relaxation and Diffusion Correlation Data. In D. Topgaard (Ed.), Advanced Diffusion Encoding Methods in MRI (pp. 0). The Royal Society of Chemistry. 10.1039/9781788019910-00278

Benjamini, D., & Basser, P. J. (2016). Use of marginal distributions constrained optimization (MADCO) for accelerated 2D MRI relaxometry and diffusometry. Journal of Magnetic Resonance, 271, 40–45. 10.1016/j.jmr.2016.08.004

Benjamini, D., & Basser, P. J. (2017). Magnetic resonance microdynamic imaging reveals distinct tissue microenvironments. NeuroImage, 163, 183–196. 10.1016/j.neuroimage.2017.09.033

Benjamini, D., & Basser, P. J. (2020). Multidimensional correlation MRI. NMR in Biomedicine, 33(12), e4226. 10.1002/nbm.4226

Benjamini, D., Bouhrara, M., Komlosh, M. E., Iacono, D., Perl, D. P., Brody, D. L., & Basser, P. J. (2021). Multidimensional MRI for Characterization of Subtle Axonal Injury Accelerated Using an Adaptive Nonlocal Multispectral Filter [Brief Research Report]. Frontiers in Physics, 9. 10.3389/fphy.2021.737374

Benjamini, D., Hutchinson, E. B., Komlosh, M. E., Comrie, C. J., Schwerin, S. C., Zhang, G., Pierpaoli, C., & Basser, P. J. (2020). Direct and specific assessment of axonal injury and spinal cord microenvironments using diffusion correlation imaging. NeuroImage, 221, 117195. 10.1016/j.neuroimage.2020.117195

Benjamini, D., Iacono, D., Komlosh, M. E., Perl, D. P., Brody, D. L., & Basser, P. J. (2021). Diffuse axonal injury has a characteristic multidimensional MRI signature in the human brain. Brain, 144(3), 800–816. 10.1093/brain/awaa447

Benjamini, D., Komlosh, M. E., Holtzclaw, L. A., Nevo, U., & Basser, P. J. (2016). White matter microstructure from nonparametric axon diameter distribution mapping. NeuroImage, 135, 333–344. 10.1016/j.neuroimage.2016.04.052

Benjamini, D., Priemer, D. S., Perl, D. P., Brody, D. L., & Basser, P. J. (2022). Mapping astrogliosis in the individual human brain using multidimensional MRI. Brain, 146(3), 1212–1226. 10.1093/brain/awac298

Bland, J. M., & Altman, D. G. (1986). Statistical methods for assessing agreement between two methods of clinical measurement. The Lancet, 327(8476), 307–310. 10.1016/S0140-6736(86)90837-8

Caceres, A., Hall, D. L., Zelaya, F. O., Williams, S. C. R., & Mehta, M. A. (2009). Measuring fMRI reliability with the intra-class correlation coefficient. NeuroImage, 45(3), 758–768. 10.1016/j.neuroimage.2008.12.035

Callaghan, P. T. (2011). Translational Dynamics and Magnetic Resonance: Principles of Pulsed Gradient Spin Echo NMR. Oxford University Press. 10.1093/acprof:oso/9780199556984.001.0001

Coelho, S., Baete, S. H., Lemberskiy, G., Ades-Aron, B., Barrol, G., Veraart, J., Novikov, D. S., & Fieremans, E. (2022). Reproducibility of the Standard Model of diffusion in white matter on clinical MRI systems. NeuroImage, 257, 119290. 10.1016/j.neuroimage.2022.119290

Conturo, T. E., McKinstry, R. C., Akbudak, E., & Robinson, B. H. (1996). Encoding of anisotropic diffusion with tetrahedral gradients: A general mathematical diffusion formalism and experimental results. Magnetic Resonance in Medicine, 35(3), 399–412. 10.1002/mrm.1910350319

Dai, E., Zhu, A., Yang, G. K., Quah, K., Tan, E. T., Fiveland, E., Foo, T. K. F., & McNab, J. A. (2023). Frequency-dependent diffusion kurtosis imaging in the human brain using an oscillating gradient spin echo sequence and a high-performance head-only gradient. NeuroImage, 279, 120328. 10.1016/j.neuroimage.2023.120328

de Almeida Martins, J. P., Tax, C. M. W., Reymbaut, A., Szczepankiewicz, F., Chamberland, M., Jones, D. K., & Topgaard, D. (2021). Computing and visualising intra-voxel orientation-specific relaxation– diffusion features in the human brain. Human Brain Mapping, 42(2), 310–328. 10.1002/hbm.25224

de Almeida Martins, J. P., Tax, C. M. W., Szczepankiewicz, F., Jones, D. K., Westin, C. F., & Topgaard, D. (2020). Transferring principles of solid-state and Laplace NMR to the field of in vivo brain MRI. Magn. Reson., 1(1), 27–43. 10.5194/mr-1-27-2020

de Almeida Martins, J. P., & Topgaard, D. (2016). Two-Dimensional Correlation of Isotropic and Directional Diffusion Using NMR. Physical Review Letters, 116(8), 087601. 10.1103/PhysRevLett.116.087601

de Almeida Martins, J. P., & Topgaard, D. (2018). Multidimensional correlation of nuclear relaxation rates and diffusion tensors for model-free investigations of heterogeneous anisotropic porous materials. Scientific Reports, 8(1), 2488. 10.1038/s41598-018-19826-9

Di Carlo, D. T., Benedetto, N., Duffau, H., Cagnazzo, F., Weiss, A., Castagna, M., Cosottini, M., & Perrini, P. (2019). Microsurgical anatomy of the sagittal stratum. Acta Neurochirurgica, 161(11), 2319–2327. 10.1007/s00701-019-04019-8

Fan, Q., Polackal, M. N., Tian, Q., Ngamsombat, C., Nummenmaa, A., Witzel, T., Klawiter, E. C., & Huang, S. Y. (2021). Scan-rescan repeatability of axonal imaging metrics using high-gradient diffusion MRI and statistical implications for study design. NeuroImage, 240, 118323. 10.1016/j.neuroimage.2021.118323

Grech-Sollars, M., Hales, P. W., Miyazaki, K., Raschke, F., Rodriguez, D., Wilson, M., Gill, S. K., Banks, T., Saunders, D. E., Clayden, J. D., Gwilliam, M. N., Barrick, T. R., Morgan, P. S., Davies, N. P., Rossiter, J., Auer, D. P., Grundy, R., Leach, M. O., Howe, F. A., … Clark, C. A. (2015). Multi-centre reproducibility of diffusion MRI parameters for clinical sequences in the brain. NMR in Biomedicine, 28(4), 468–485. 10.1002/nbm.3269

Hagiwara, A., Hori, M., Cohen-Adad, J., Nakazawa, M., Suzuki, Y., Kasahara, A., Horita, M., Haruyama, T., Andica, C., Maekawa, T., Kamagata, K., Kumamaru, K. K., Abe, O., & Aoki, S. (2019). Linearity, Bias, Intrascanner Repeatability, and Interscanner Reproducibility of Quantitative Multidynamic Multiecho Sequence for Rapid Simultaneous Relaxometry at 3 T: A Validation Study With a Standardized Phantom and Healthy Controls. Investigative Radiology, 54(1), 39–47. 10.1097/rli.0000000000000510

Hanspach, J., Nagel, A. M., Hensel, B., Uder, M., Koros, L., & Laun, F. B. (2021). Sample size estimation: Current practice and considerations for original investigations in MRI technical development studies. Magnetic Resonance in Medicine, 85(4), 2109–2116. 10.1002/mrm.28550

Huang, S. Y., Witzel, T., Keil, B., Scholz, A., Davids, M., Dietz, P., Rummert, E., Ramb, R., Kirsch, J. E., Yendiki, A., Fan, Q., Tian, Q., Ramos-Llordén, G., Lee, H.-H., Nummenmaa, A., Bilgic, B., Setsompop, K., Wang, F., Avram, A. V., Rosen, B. R. (2021). Connectome 2.0: Developing the next-generation ultra-high gradient strength human MRI scanner for bridging studies of the micro-, meso- and macro-connectome. NeuroImage, 243, 118530. 10.1016/j.neuroimage.2021.118530

Huo, Y., Xu, Z., Xiong, Y., Aboud, K., Parvathaneni, P., Bao, S., Bermudez, C., Resnick, S. M., Cutting, L. E., & Landman, B. A. (2019). 3D whole brain segmentation using spatially localized atlas network tiles. NeuroImage, 194, 105–119. 10.1016/j.neuroimage.2019.03.041

Hürlimann, M. D., & Venkataramanan, L. (2002). Quantitative Measurement of Two-Dimensional Distribution Functions of Diffusion and Relaxation in Grossly Inhomogeneous Fields. Journal of Magnetic Resonance, 157(1), 31–42. 10.1006/jmre.2002.2567

Irfanoglu, M. O., Modi, P., Nayak, A., Hutchinson, E. B., Sarlls, J., & Pierpaoli, C. (2015). DR-BUDDI (Diffeomorphic Registration for Blip-Up blip-Down Diffusion Imaging) method for correcting echo planar imaging distortions. NeuroImage, 106, 284–299. 10.1016/j.neuroimage.2014.11.042

Jespersen, S. N., Olesen, J. L., Ianuş, A., & Shemesh, N. (2019). Effects of nongaussian diffusion on “isotropic diffusion” measurements: An ex-vivo microimaging and simulation study. Journal of Magnetic Resonance, 300, 84–94. 10.1016/j.jmr.2019.01.007

Johnson, J. T. E., Irfanoglu, M. O., Manninen, E., Ross, T. J., Yang, Y., Laun, F. B., Martin, J., Topgaard, D., & Benjamini, D. (2024). In vivo disentanglement of diffusion frequency-dependence, tensor shape, and relaxation using multidimensional MRI. Human Brain Mapping. 10.1002/hbm.26697

Kellner, E., Dhital, B., Kiselev, V. G., & Reisert, M. (2016). Gibbs-ringing artifact removal based on local subvoxel-shifts. Magnetic Resonance in Medicine, 76(5), 1574–1581. 10.1002/mrm.26054

Kim, D., Doyle, E. K., Wisnowski, J. L., Kim, J. H., & Haldar, J. P. (2017). Diffusion-relaxation correlation spectroscopic imaging: A multidimensional approach for probing microstructure. Magnetic Resonance in Medicine, 78(6), 2236–2249. 10.1002/mrm.26629

Koo, T. K., & Li, M. Y. (2016). A Guideline of Selecting and Reporting Intraclass Correlation Coefficients for Reliability Research. Journal of Chiropractic Medicine, 15(2), 155–163. 10.1016/j.jcm.2016.02.012

Kundu, S., Barsoum, S., Ariza, J., Nolan, A. L., Latimer, C. S., Keene, C. D., Basser, P. J., & Benjamini, D. (2023). Mapping the individual human cortex using multidimensional MRI and unsupervised learning. Brain Communications, 5(6). 10.1093/braincomms/fcad258

Lampinen, B., Szczepankiewicz, F., Novén, M., van Westen, D., Hansson, O., Englund, E., Mårtensson, J., Westin, C.-F., & Nilsson, M. (2019). Searching for the neurite density with diffusion MRI: Challenges for biophysical modeling. Human Brain Mapping, 40(8), 2529–2545. 10.1002/hbm.24542

Lasič, S., Szczepankiewicz, F., Eriksson, S., Nilsson, M., & Topgaard, D. (2014). Microanisotropy imaging: quantification of microscopic diffusion anisotropy and orientational order parameter by diffusion MRI with magic-angle spinning of the q-vector [Original Research]. Frontiers in Physics, 2. 10.3389/fphy.2014.00011

Lasič, S., Yuldasheva, N., Szczepankiewicz, F., Nilsson, M., Budde, M., Dall’Armellina, E., Schneider, J. E., Teh, I., & Lundell, H. (2022). Stay on the Beat With Tensor-Valued Encoding: Time-Dependent Diffusion and Cell Size Estimation in ex vivo Heart [Original Research]. Frontiers in Physics, 10. 10.3389/fphy.2022.812115

Lawrenz, M., Koch, M. A., & Finsterbusch, J. (2010). A tensor model and measures of microscopic anisotropy for double-wave-vector diffusion-weighting experiments with long mixing times. Journal of Magnetic Resonance, 202(1), 43–56. 10.1016/j.jmr.2009.09.015

Lee, H.-H., Novikov, D. S., & Fieremans, E. (2021). Removal of partial Fourier-induced Gibbs (RPG) ringing artifacts in MRI. Magnetic Resonance in Medicine, 86(5), 2733–2750. 10.1002/mrm.28830

Livet, J., Weissman, T. A., Kang, H., Draft, R. W., Lu, J., Bennis, R. A., Sanes, J. R., & Lichtman, J. W. (2007). Transgenic strategies for combinatorial expression of fluorescent proteins in the nervous system. Nature, 450(7166), 56–62. 10.1038/nature06293

Lundell, H., & Lasič, S. (2020). Diffusion Encoding with General Gradient Waveforms. In D. Topgaard (Ed.), Advanced Diffusion Encoding Methods in MRI (pp. 0). The Royal Society of Chemistry. 10.1039/9781788019910-00012

Luque Laguna, P. A., Combes, A. J. E., Streffer, J., Einstein, S., Timmers, M., Williams, S. C. R., & Dell’Acqua, F. (2020). Reproducibility, reliability and variability of FA and MD in the older healthy population: A test-retest multiparametric analysis. NeuroImage: Clinical, 26, 102168. 10.1016/j.nicl.2020.102168

Mackay, A., Whittall, K., Adler, J., Li, D., Paty, D., & Graeb, D. (1994). In vivo visualization of myelin water in brain by magnetic resonance. Magnetic Resonance in Medicine, 31(6), 673–677. 10.1002/mrm.1910310614

Manning, A. P., MacKay, A. L., & Michal, C. A. (2021). Understanding aqueous and non-aqueous proton T1 relaxation in brain. Journal of Magnetic Resonance, 323, 106909. 10.1016/j.jmr.2020.106909

Martin, J., Reymbaut, A., Schmidt, M., Doerfler, A., Uder, M., Laun, F. B., & Topgaard, D. (2021). Nonparametric D-R1-R2 distribution MRI of the living human brain. NeuroImage, 245, 118753. 10.1016/j.neuroimage.2021.118753

Matheson, G. J. (2019). We need to talk about reliability: making better use of test-retest studies for study design and interpretation. PeerJ, 7, e6918. 10.7717/peerj.6918

McGraw, K. O., & Wong, S. P. (1996). Forming inferences about some intraclass correlation coefficients. Psychological Methods, 1(1), 30–46. 10.1037/1082-989X.1.1.30

Morel, A. (2007). Stereotactic Atlas of the Human Thalamus and Basal Ganglia (1 ed.). CRC Press. 10.3109/9781420016796

Naranjo, I. D., Reymbaut, A., Brynolfsson, P., Lo Gullo, R., Bryskhe, K., Topgaard, D., Giri, D. D., Reiner, J. S., Thakur, S. B., & Pinker-Domenig, K. (2021). Multidimensional Diffusion Magnetic Resonance Imaging for Characterization of Tissue Microstructure in Breast Cancer Patients: A Prospective Pilot Study. Cancers, 13(7), 1606. 10.3390/cancers13071606

Narvaez, O., Svenningsson, L., Yon, M., Sierra, A., & Topgaard, D. (2022). Massively Multidimensional Diffusion-Relaxation Correlation MRI [Original Research]. Frontiers in Physics, 9. 10.3389/fphy.2021.793966

Neuman, C. H. (1974). Spin echo of spins diffusing in a bounded medium. The Journal of Chemical Physics, 60(11), 4508–4511. 10.1063/1.1680931

Nilsson, M., Szczepankiewicz, F., Lampinen, B., Ahlgren, A., de Almeida Martins, J. P., Lasic, S., Westin, C.-F., & Topgaard, D. (2018). An open-source framework for analysis of multidimensional diffusion MRI data implemented in MATLAB.Proc. Intl. Soc. Mag. Reson. Med. Paris, France.

Novello, L., Henriques, R. N., Ianuş, A., Feiweier, T., Shemesh, N., & Jovicich, J. (2022). In vivo Correlation Tensor MRI reveals microscopic kurtosis in the human brain on a clinical 3T scanner. NeuroImage, 254, 119137. 10.1016/j.neuroimage.2022.119137

Novikov, D. S., Fieremans, E., Jespersen, S. N., & Kiselev, V. G. (2019). Quantifying brain microstructure with diffusion MRI: Theory and parameter estimation. NMR in Biomedicine, 32(4), e3998. 10.1002/nbm.3998

Novikov, D. S., Kiselev, V. G., & Jespersen, S. N. (2018). On modeling. Magnetic Resonance in Medicine, 79(6), 3172–3193. 10.1002/mrm.27101

Peled, S., Cory, D. G., Raymond, S. A., Kirschner, D. A., & Jolesz, F. A. (1999). Water diffusion, T2, and compartmentation in frog sciatic nerve. Magnetic Resonance in Medicine, 42(5), 911–918. 10.1002/(SICI)1522-2594(199911)42:5<911::AID-MRM11>3.0.CO;2-J

Pierpaoli, C., Jezzard, P., Basser, P. J., Barnett, A., & Chiro, G. D. (1996). Diffusion tensor MR imaging of the human brain. Radiology, 201(3), 637–648. 10.1148/radiology.201.3.8939209

Reuter, M., Schmansky, N. J., Rosas, H. D., & Fischl, B. (2012). Within-subject template estimation for unbiased longitudinal image analysis. NeuroImage, 61(4), 1402–1418. 10.1016/j.neuroimage.2012.02.084

Reymbaut, A., Critchley, J., Durighel, G., Sprenger, T., Sughrue, M., Bryskhe, K., & Topgaard, D. (2021). Toward nonparametric diffusion-characterization of crossing fibers in the human brain. Magnetic Resonance in Medicine, 85(5), 2815–2827. 10.1002/mrm.28604

Reymbaut, A., Mezzani, P., de Almeida Martins, J. P., & Topgaard, D. (2020). Accuracy and precision of statistical descriptors obtained from multidimensional diffusion signal inversion algorithms. NMR in Biomedicine, 33(12), e4267. 10.1002/nbm.4267

Rohde, G. K., Barnett, A. S., Basser, P. J., Marenco, S., & Pierpaoli, C. (2004). Comprehensive approach for correction of motion and distortion in diffusion-weighted MRI. Magnetic Resonance in Medicine, 51(1), 103–114. 10.1002/mrm.10677

Schilling, K. G., Blaber, J., Huo, Y., Newton, A., Hansen, C., Nath, V., Shafer, A. T., Williams, O., Resnick, S. M., Rogers, B., Anderson, A. W., & Landman, B. A. (2019). Synthesized b0 for diffusion distortion correction (Synb0-DisCo). Magnetic Resonance Imaging, 64, 62–70. 10.1016/j.mri.2019.05.008

Shemesh, N., Jespersen, S. N., Alexander, D. C., Cohen, Y., Drobnjak, I., Dyrby, T. B., Finsterbusch, J., Koch, M. A., Kuder, T., Laun, F., Lawrenz, M., Lundell, H., Mitra, P. P., Nilsson, M., Özarslan, E., Topgaard, D., & Westin, C.-F. (2016). Conventions and nomenclature for double diffusion encoding NMR and MRI. Magnetic Resonance in Medicine, 75(1), 82–87. 10.1002/mrm.25901

Silva, M. D., Helmer, K. G., Lee, J.-H., Han, S. S., Springer, C. S., & Sotak, C. H. (2002). Deconvolution of Compartmental Water Diffusion Coefficients in Yeast-Cell Suspensions Using Combined T1 and Diffusion Measurements. Journal of Magnetic Resonance, 156(1), 52–63. 10.1006/jmre.2002.2527

Sjölund, J., Szczepankiewicz, F., Nilsson, M., Topgaard, D., Westin, C.-F., & Knutsson, H. (2015). Constrained optimization of gradient waveforms for generalized diffusion encoding. Journal of Magnetic Resonance, 261, 157–168. 10.1016/j.jmr.2015.10.012

Slator, P. J., Hutter, J., Palombo, M., Jackson, L. H., Ho, A., Panagiotaki, E., Chappell, L. C., Rutherford, M. A., Hajnal, J. V., & Alexander, D. C. (2019). Combined diffusion-relaxometry MRI to identify dysfunction in the human placenta. Magnetic Resonance in Medicine, 82(1), 95–106. 10.1002/mrm.27733

Slator, P. J., Palombo, M., Miller, K. L., Westin, C.-F., Laun, F., Kim, D., Haldar, J. P., Benjamini, D., Lemberskiy, G., de Almeida Martins, J. P., & Hutter, J. (2021). Combined diffusion-relaxometry microstructure imaging: Current status and future prospects. Magnetic Resonance in Medicine, 86(6), 2987–3011. 10.1002/mrm.28963

Soares, J., Marques, P., Alves, V., & Sousa, N. (2013). A hitchhiker’s guide to diffusion tensor imaging [Review]. Frontiers in Neuroscience, 7. 10.3389/fnins.2013.00031

Stanisz, G. J., & Henkelman, R. M. (1998). Diffusional anisotropy of T2 components in bovine optic nerve. Magnetic Resonance in Medicine, 40(3), 405–410. 10.1002/mrm.1910400310

Stanisz, G. J., Wright, G. A., Henkelman, R. M., & Szafer, A. (1997). An analytical model of restricted diffusion in bovine optic nerve. Magnetic Resonance in Medicine, 37(1), 103–111. 10.1002/mrm.1910370115

Stepišnik, J. (1981). Analysis of NMR self-diffusion measurements by a density matrix calculation. Physica B+C, 104(3), 350–364. 10.1016/0378-4363(81)90182-0

Stepišnik, J. (1993). Time-dependent self-diffusion by NMR spin-echo. Physica B: Condensed Matter, 183(4), 343–350. 10.1016/0921-4526(93)90124-O

Tofts, P. (2003). Quantitative MRI of the Brain: Measuring Changes Caused by Disease. John Wiley & Sons, Ltd. 10.1002/0470869526

Topgaard, D. (2019). Diffusion tensor distribution imaging. NMR in Biomedicine, 32(5), e4066. 10.1002/nbm.4066

Vasilescu, V., Katona, E., Simplâceanu, V., & Demco, D. (1978). Water compartments in the myelinated nerve. III. Pulsed NMR result. Experientia, 34(11), 1443–1444. 10.1007/BF01932339

Veraart, J., Fieremans, E., & Novikov, D. S. (2019). On the scaling behavior of water diffusion in human brain white matter. NeuroImage, 185, 379–387. 10.1016/j.neuroimage.2018.09.075

Veraart, J., Novikov, D. S., Christiaens, D., Ades-aron, B., Sijbers, J., & Fieremans, E. (2016). Denoising of diffusion MRI using random matrix theory. NeuroImage, 142, 394–406. 10.1016/j.neuroimage.2016.08.016

Veraart, J., Nunes, D., Rudrapatna, U., Fieremans, E., Jones, D. K., Novikov, D. S., & Shemesh, N. (2020). Noninvasive quantification of axon radii using diffusion MRI. eLife, 9, e49855. 10.7554/eLife.49855

Wei, X., Zhu, L., Zeng, Y., Xue, K., Dai, Y., Xu, J., Liu, G., Liu, F., Xue, W., Wu, D., & Wu, G. (2022). Detection of prostate cancer using diffusion-relaxation correlation spectrum imaging with support vector machine model – a feasibility study. Cancer Imaging, 22(1), 77. 10.1186/s40644-022-00516-9

Weiskopf, N., Edwards, L. J., Helms, G., Mohammadi, S., & Kirilina, E. (2021). Quantitative magnetic resonance imaging of brain anatomy and in vivo histology. Nature Reviews Physics, 3(8), 570–588. 10.1038/s42254-021-00326-1

Wetscherek, A., Stieltjes, B., & Laun, F. B. (2015). Flow-compensated intravoxel incoherent motion diffusion imaging. Magnetic Resonance in Medicine, 74(2), 410–419. 10.1002/mrm.25410

Woessner, D. E. (1963). N.M.R. spin-echo self-diffusion measurements on fluids undergoing restricted diffusion. The Journal of Physical Chemistry, 67(6), 1365–1367. 10.1021/j100800a509

Xiong, Y., Huo, Y., Wang, J., Davis, L. T., McHugo, M., & Landman, B. (2019). Reproducibility evaluation of SLANT whole brain segmentation across clinical magnetic resonance imaging protocols (Vol. 10949). SPIE. 10.1117/12.2512561

Yon, M., de Almeida Martins, J. P., Bao, Q., Budde, M. D., Frydman, L., & Topgaard, D. (2020). Diffusion tensor distribution imaging of an in vivo mouse brain at ultrahigh magnetic field by spatiotemporal encoding. NMR in Biomedicine, 33(11), e4355. 10.1002/nbm.4355

Zhang, H., Schneider, T., Wheeler-Kingshott, C. A., & Alexander, D. C. (2012). NODDI: Practical in vivo neurite orientation dispersion and density imaging of the human brain. NeuroImage, 61(4), 1000–1016. 10.1016/j.neuroimage.2012.03.072

