## Supplementary Fig for "The variability of multidimensional diffusion-relaxation MRI estimates in the human brain"

### Supplementary Figures

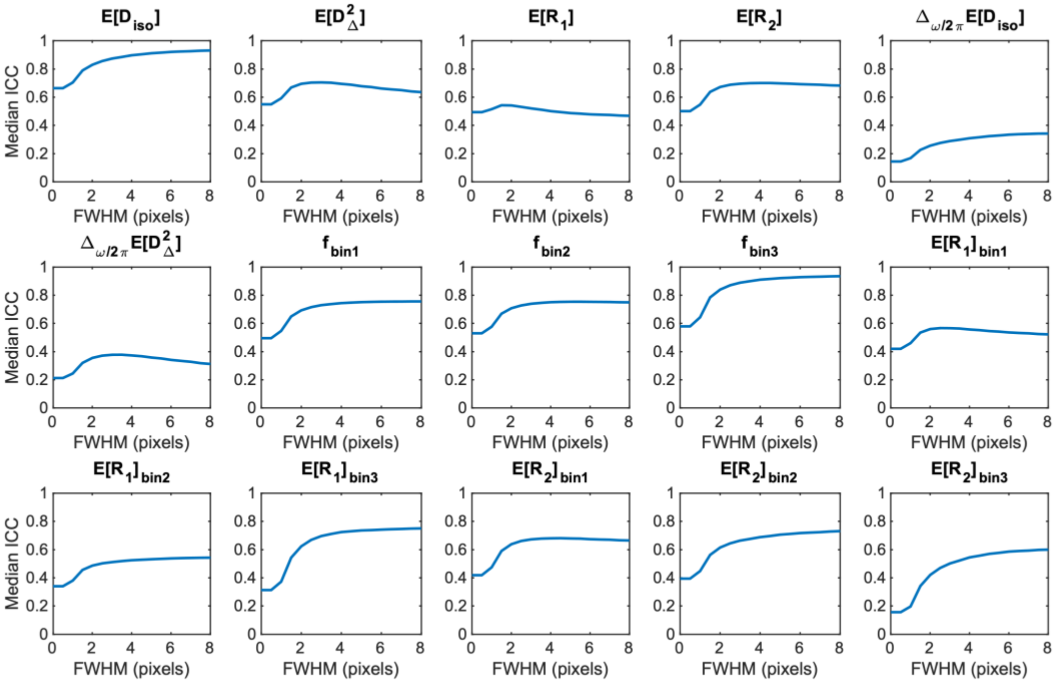


Supplementary Fig. 1. The dependence of median voxel-wise intraclass correlation coefficient (ICC; reliability) on spatial smoothing. The median ICC was computed over all brain voxels in the cohort template space at different values of the full width at half maximum (FWHM) of the Gaussian spatial smoothing kernel. One pixel corresponds to 2 mm.

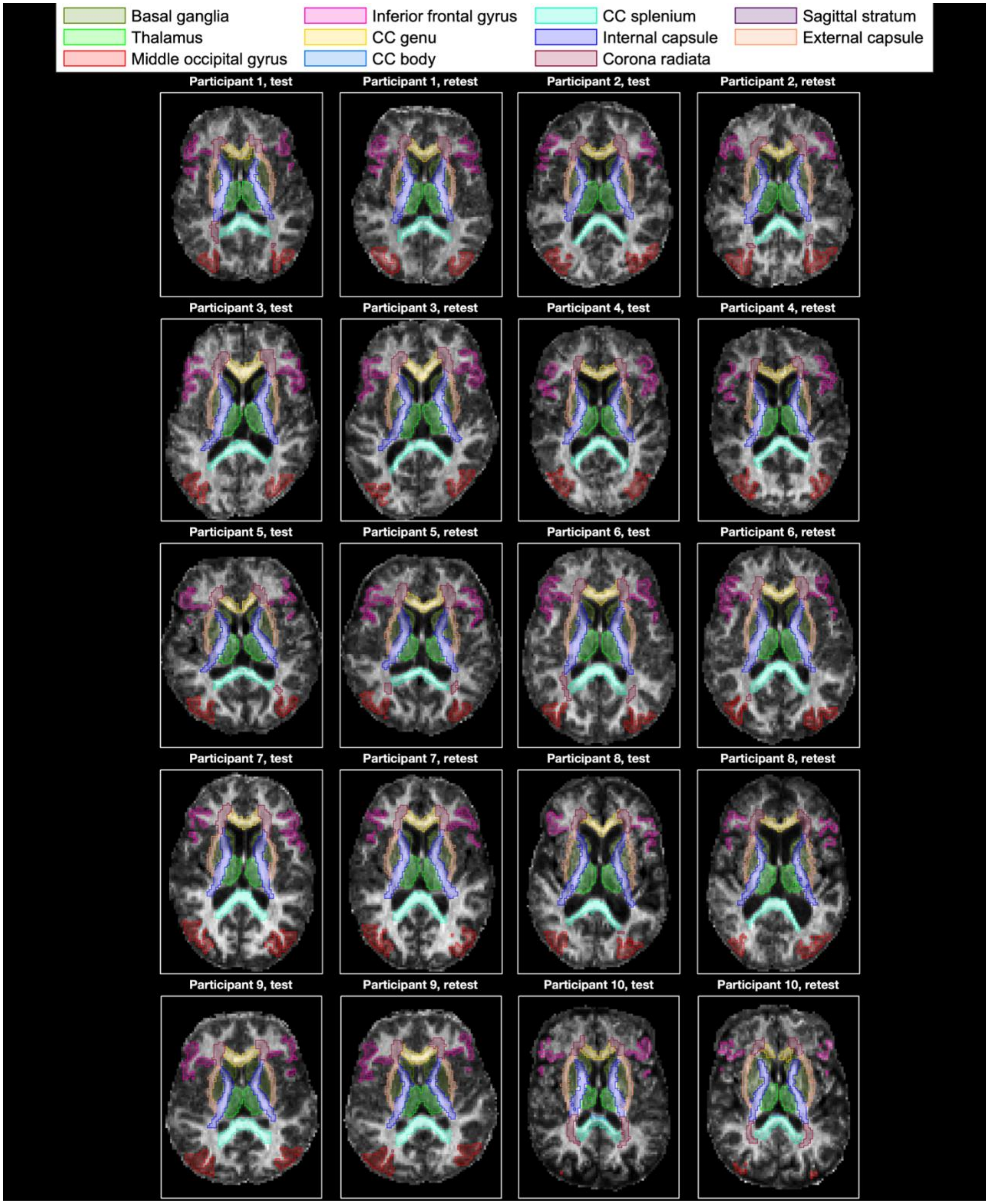


Supplementary Fig. 2. Regions of interest (ROI) in the native space shown for each repetition and participant. The ROIs are overlaid on top of the $\mathrm{E}[D_{\Delta}^{2}]$ map.

**A**

**Mean**

| | Basal ganglia | Thalamus | Middle occipital gyrus | Inferior frontal gyrus | CC genu | CC body | CC splenium | Internal capsule | Corona radiata | Sagittal stratum | External capsule |
|---|---|---|---|---|---|---|---|---|---|---|---|
| $E[D_{iso}]$ (μm²/ms) | 0.93 | 1.01 | 0.95 | 1.05 | 1.11 | 1.10 | 1.09 | 1.03 | 1.06 | 1.15 | 1.01 |
| $E[D_{\Delta}^{2}]$ | 0.32 | 0.38 | 0.27 | 0.25 | 0.57 | 0.59 | 0.67 | 0.62 | 0.59 | 0.52 | 0.46 |
| $E[R_1]$ ($s^{-1}$) | 0.92 | 0.99 | 0.76 | 0.71 | 1.07 | 1.05 | 1.13 | 1.08 | 1.06 | 1.15 | 0.97 |
| $E[R_2]$ ($s^{-1}$) | 20.70 | 17.90 | 16.60 | 15.20 | 17.00 | 15.60 | 16.70 | 17.50 | 15.50 | 16.60 | 16.70 |
| $\Delta_{\omega/2\pi}E[D_{iso}]$ (μm²) | 1.30 | 1.00 | 6.90 | 0.80 | -0.40 | 0.90 | 1.20 | 1.10 | 1.50 | -0.20 | -0.10 |
| $\Delta_{\omega/2\pi}E[D_{\Delta}^{2}]$ (ms) | -2.01 | -2.50 | -2.20 | -2.00 | -2.80 | -3.20 | -2.40 | -2.80 | -3.80 | -2.70 | -2.40 |
| $f_{bin1}$ | 0.41 | 0.51 | 0.36 | 0.31 | 0.72 | 0.74 | 0.84 | 0.80 | 0.76 | 0.65 | 0.60 |
| $f_{bin2}$ | 0.56 | 0.46 | 0.60 | 0.62 | 0.23 | 0.21 | 0.12 | 0.17 | 0.21 | 0.30 | 0.37 |
| $f_{bin3}$ | 0.01 | 0.01 | 0.03 | 0.05 | 0.02 | 0.02 | 0.01 | 0.00 | 0.00 | 0.01 | 0.01 |
| $E[R_1]_{bin1}$ ($s^{-1}$) | 0.82 | 0.96 | 0.63 | 0.58 | 1.09 | 1.06 | 1.15 | 1.09 | 1.07 | 1.16 | 0.98 |
| $E[R_1]_{bin2}$ ($s^{-1}$) | 0.88 | 0.94 | 0.78 | 0.74 | 0.66 | 0.62 | 0.41 | 0.51 | 0.70 | 1.03 | 0.84 |
| $E[R_1]_{bin3}$ ($s^{-1}$) | 0.02 | 0.03 | 0.06 | 0.11 | 0.05 | 0.05 | 0.04 | 0.01 | 0.01 | 0.05 | 0.02 |
| $E[R_2]_{bin1}$ ($s^{-1}$) | 20.60 | 17.90 | 16.30 | 15.00 | 16.70 | 14.60 | 16.10 | 17.00 | 15.10 | 15.50 | 16.90 |
| $E[R_2]_{bin2}$ ($s^{-1}$) | 18.40 | 15.00 | 15.10 | 14.30 | 10.40 | 9.60 | 5.70 | 8.50 | 9.60 | 14.10 | 13.10 |
| $E[R_2]_{bin3}$ ($s^{-1}$) | 0.50 | 0.70 | 0.80 | 1.40 | 1.00 | 1.00 | 0.80 | 0.30 | 0.10 | 1.00 | 0.40 |

**B**

**SD**

| | Basal ganglia | Thalamus | Middle occipital gyrus | Inferior frontal gyrus | CC genu | CC body | CC splenium | Internal capsule | Corona radiata | Sagittal stratum | External capsule |
|---|---|---|---|---|---|---|---|---|---|---|---|
| $E[D_{iso}]$ (μm²/ms) | 0.19 | 0.17 | 0.20 | 0.22 | 0.16 | 0.17 | 0.17 | 0.14 | 0.10 | 0.14 | 0.15 |
| $E[D_{\Delta}^{2}]$ | 0.12 | 0.12 | 0.11 | 0.08 | 0.13 | 0.14 | 0.15 | 0.15 | 0.10 | 0.08 | 0.12 |
| $E[R_1]$ ($s^{-1}$) | 0.12 | 0.12 | 0.13 | 0.12 | 0.15 | 0.15 | 0.14 | 0.13 | 0.10 | 0.13 | 0.13 |
| $E[R_2]$ ($s^{-1}$) | 3.70 | 2.30 | 2.30 | 2.10 | 2.30 | 2.20 | 2.30 | 2.70 | 1.90 | 1.80 | 2.10 |
| $\Delta_{\omega/2\pi}E[D_{iso}]$ (μm²) | 4.70 | 5.30 | 5.90 | 4.80 | 4.30 | 4.20 | 3.30 | 3.50 | 3.80 | 4.40 | 4.40 |
| $\Delta_{\omega/2\pi}E[D_{\Delta}^{2}]$ (ms) | 0.99 | 1.20 | 1.20 | 1.10 | 1.50 | 1.90 | 1.60 | 1.50 | 1.80 | 1.30 | 1.20 |
| $f_{bin1}$ | 0.19 | 0.18 | 0.18 | 0.15 | 0.19 | 0.20 | 0.18 | 0.20 | 0.16 | 0.13 | 0.18 |
| $f_{bin2}$ | 0.20 | 0.18 | 0.16 | 0.14 | 0.18 | 0.19 | 0.16 | 0.19 | 0.15 | 0.13 | 0.18 |
| $f_{bin3}$ | 0.04 | 0.04 | 0.06 | 0.07 | 0.04 | 0.04 | 0.04 | 0.02 | 0.01 | 0.03 | 0.03 |
| $E[R_1]_{bin1}$ ($s^{-1}$) | 0.22 | 0.17 | 0.25 | 0.23 | 0.17 | 0.16 | 0.14 | 0.14 | 0.11 | 0.14 | 0.17 |
| $E[R_1]_{bin2}$ ($s^{-1}$) | 0.18 | 0.21 | 0.13 | 0.12 | 0.44 | 0.47 | 0.49 | 0.47 | 0.43 | 0.29 | 0.25 |
| $E[R_1]_{bin3}$ ($s^{-1}$) | 0.07 | 0.12 | 0.13 | 0.16 | 0.15 | 0.17 | 0.15 | 0.06 | 0.06 | 0.17 | 0.08 |
| $E[R_2]_{bin1}$ ($s^{-1}$) | 4.60 | 2.60 | 4.70 | 4.40 | 2.30 | 2.30 | 2.40 | 2.90 | 2.10 | 2.00 | 2.30 |
| $E[R_2]_{bin2}$ ($s^{-1}$) | 4.50 | 4.00 | 2.00 | 1.90 | 6.60 | 7.10 | 6.90 | 7.80 | 5.90 | 4.10 | 4.00 |
| $E[R_2]_{bin3}$ ($s^{-1}$) | 2.60 | 2.90 | 2.60 | 3.00 | 3.40 | 3.30 | 2.90 | 1.80 | 1.30 | 3.50 | 2.00 |

Supplementary Fig. 3. (A) Mean and (B) standard deviation (SD) of each MD-MRI parameter map, computed over all participants, repetitions, and voxels for each region of interest (ROI). Color scaling is normalized separately for each parameter map (row; darker blue is higher) to enable comparison between ROIs for each map. Abbreviations: CC, corpus callosum.

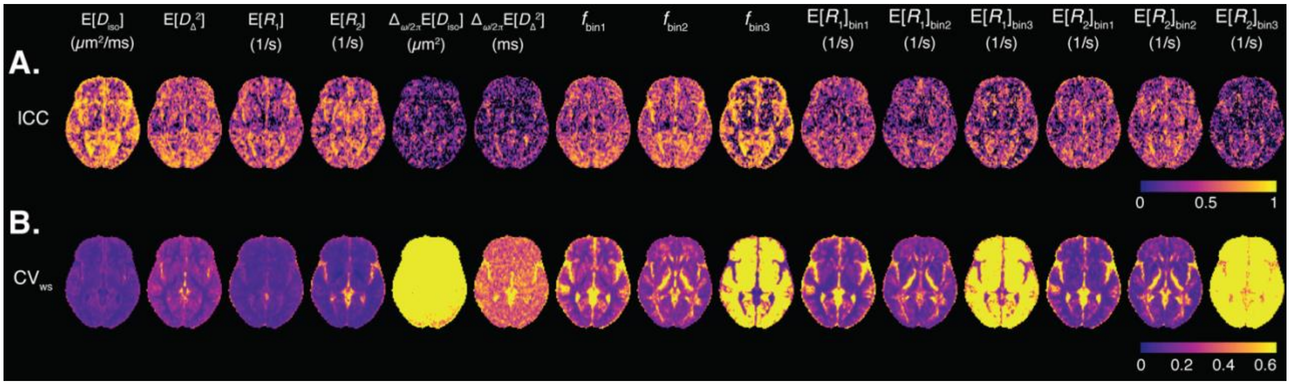


Supplementary Fig. 4. Voxel-wise intraclass correlation coefficient (ICC; reliability) and within-subject coefficient of variation ($CV_{ws}$; repeatability) without the application of spatial filtering to the maps.

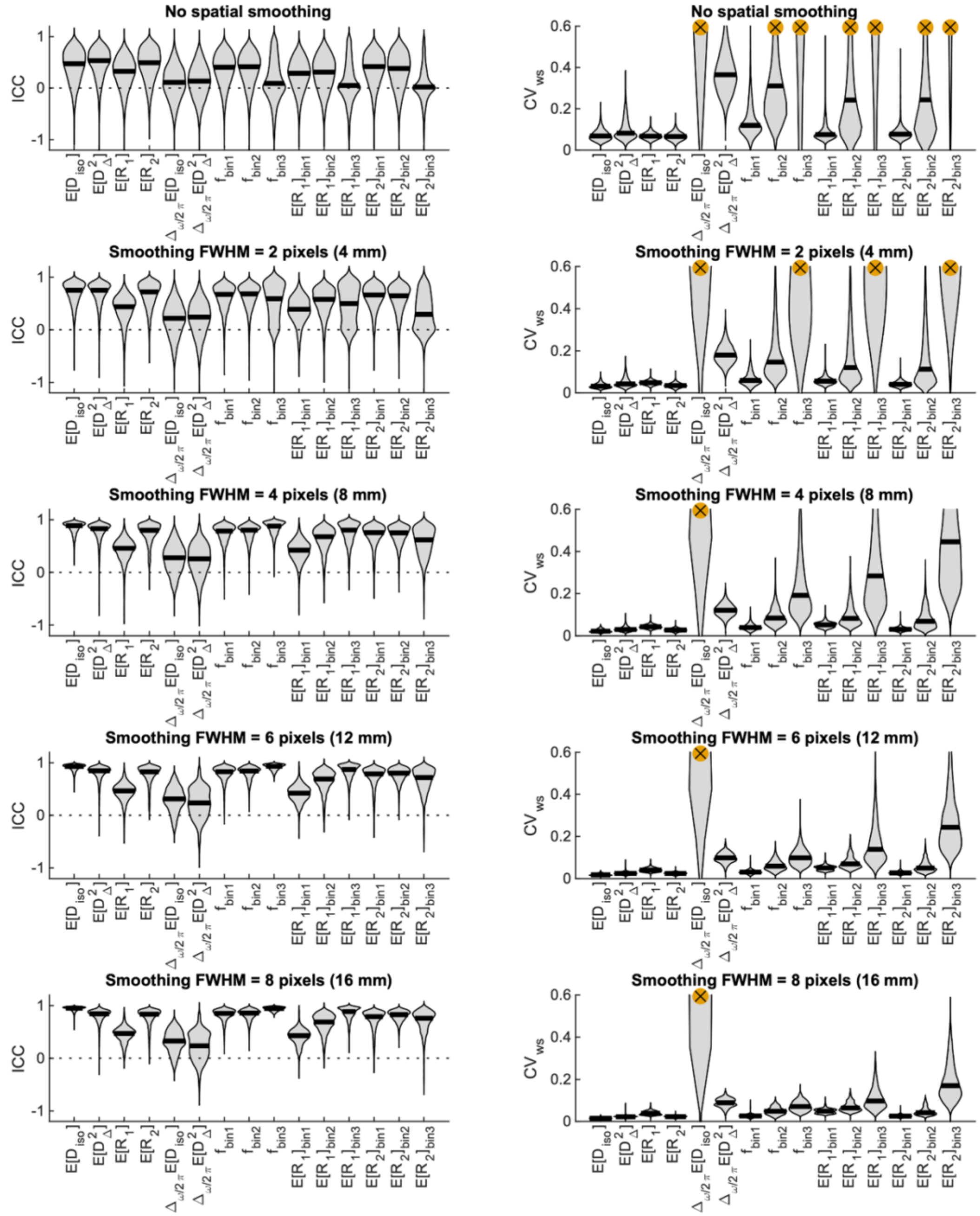


Supplementary Fig. 5. Distributions and medians of voxel-wise reliability and repeatability measures of MD-MRI parameter maps over the whole brain. The results are presented with different levels of spatial filtering (Gaussian kernel) applied to the voxel-wise parameter maps before the computation of the reliability and repeatability measures. (A) Intraclass correlation coefficient (ICC; reliability). (B) Within-subject coefficient of variation ($CV_{ws}$; repeatability). Some of the distributions included severe outliers that hindered the visualization. Such outliers were removed, and the affected distributions were marked with crosses (see main text).

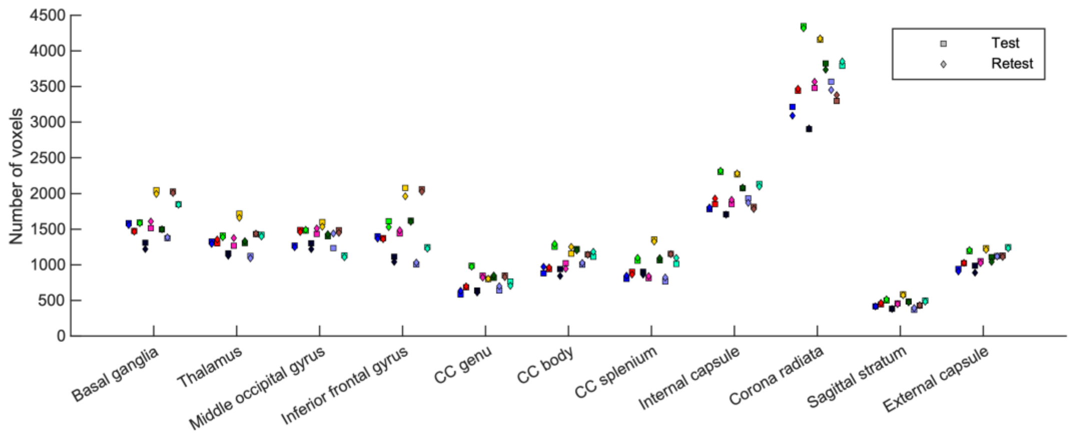


Supplementary Fig. 6. Number of voxels for each region of interest (ROI), participant, and repetition. Different colors represent different participants. Repetition 1 (test) is indicated with a square and repetition 2 (retest) with a diamond marker. Abbreviations: CC, corpus callosum.

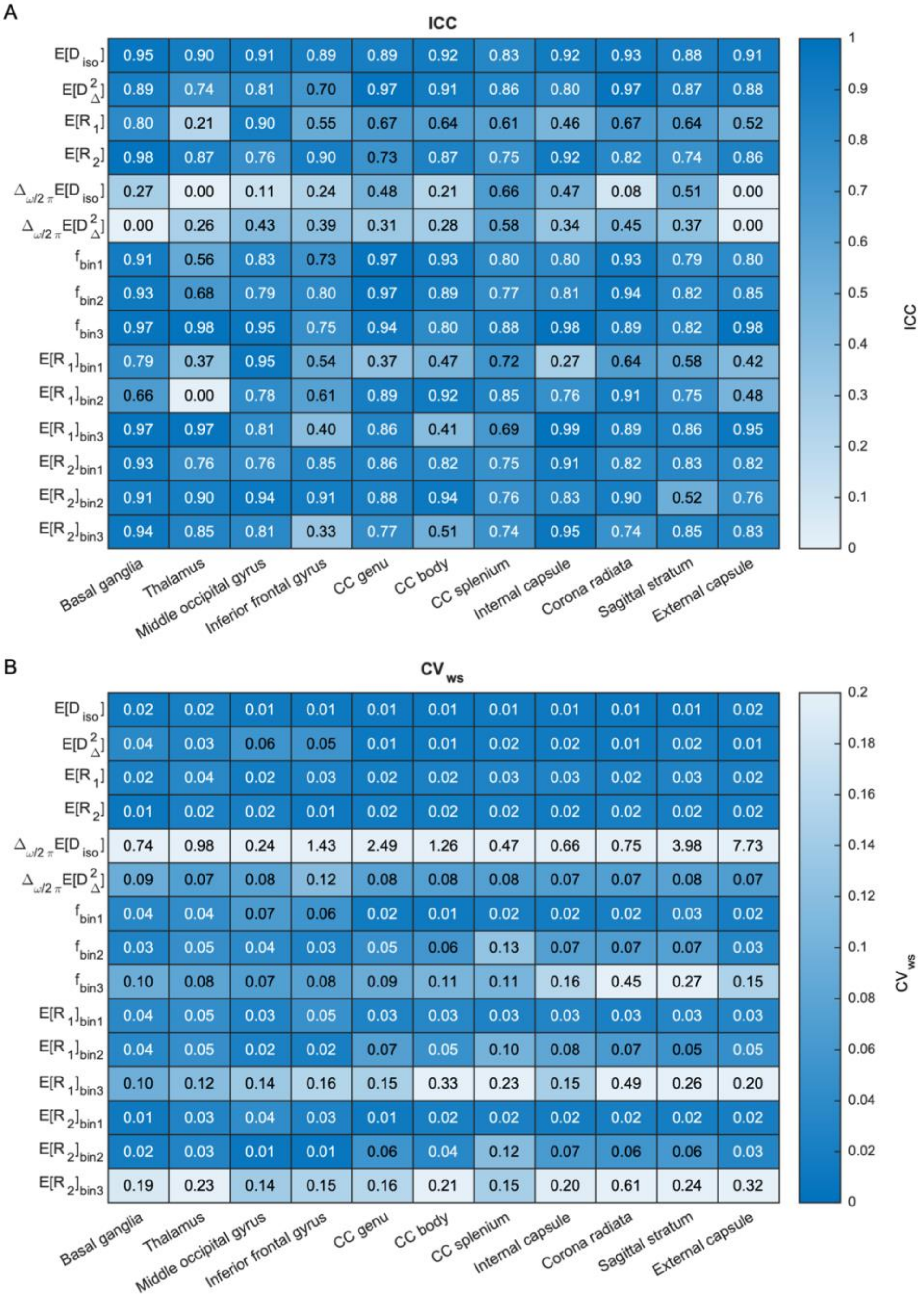


Supplementary Fig. 7. Reliability and repeatability of region-of-interest-averaged MD-MRI parameter maps. (A) Intraclass correlation coefficient (ICC; reliability). (B) Within-subject coefficient of variation ($CV_{ws}$; repeatability). Generally, ROI-based ICC values were higher and $CV_{ws}$ values lower than those of individual voxels (Figures 1 and 2). That was expected due to the increase in signal-to-noise ratio achieved through averaging. Abbreviations: CC, corpus callosum.
